# Historical ecology and stakeholder perspectives can inform peatland fire management

**DOI:** 10.64898/2026.02.19.706730

**Authors:** Jessie Woodbridge, Gina Kallis, Laura Scoble, Francis Rowney, Claire Kelly, Althea Davies

## Abstract

Climate change is increasing wildfire risk globally and peatlands have become increasingly vulnerable to fire in recent decades. We combine social research methods with analysis of historical ecological (palaeoecological) records to understand links between fire, climate, vegetation and land use. Stakeholders in the Peak District (UK) highlight the need for scientific research and local knowledge to be more frequently embedded into policy. Analyses of historical ecological datasets reveal regime shifts in moorland vegetation following periods of fire activity and managed burning. This type of disturbance can lead to dominance of grasses, which may have a negative impact on peatland carbon balance under warmer climatic conditions. Recent fires are contributing to the loss of *Sphagnum* moss and greater dominance of heath and grass. Restoring peatlands, through re-establishing native woodland and improving peat bog hydrological conditions, alongside careful planning around controlled burning, are key measures to enhance resilience to future fire events.

## Introduction

### Fire and peatlands

Rising temperatures, prolonged dry periods, and fuel accumulation are factors leading to enhanced vulnerability to wildfire globally (IPCC, 2022). Long-term climate change impacts and disturbance associated with human activity have already been observed and are projected to increase in high-carbon peatland ecosystems (Swindles et al., 2019; Sim et al., 2023). With most wildfires identified as being initiated by human action, this highlights the need for greater intervention to enhance the resilience of peatland ecosystems (Wilkinson et al., 2022). Wildfires can have major impacts on the carbon storage capacity of peat, ecosystem function, water security and biodiversity, with social and economic consequences (Baker et al., 2025). The management of wildfire and controlled burning is complex within UK upland landscapes (Harper et al., 2018) and protecting peatland carbon stocks has been identified as key to achieving emissions reduction targets (Glaves, et al., 2020). Water level changes play a crucial role in determining fire intensity in peatlands (Kartiwa et al., 2023), as fluctuations in water levels directly influence peat moisture content, which in turn affects the likelihood, spread, and severity of peat fires.

Fire is increasingly recognised as a socio-ecological process driven by complex interactions between biophysical and socio-economic factors (Perkins et al., 2022). These interacting factors need to be understood to support current and future fire management planning. Resilience can be defined as the capacity of a system to absorb disturbance and still retain the same function and structure (Walker et al., 2004). Understanding the factors that make socio-ecological systems resilient to change (Kelly et al., 2015) is key to recognising how peatlands have responded in the past and may respond to future shifts in management, climate or land use (Dougill et al., 2006). This research aims to integrate information about long-term historic ecological trends in response to fire with knowledge of recent fire risk and management. The objectives of this study are (1) to identify the key questions stakeholders have regarding peatland fires and (2) to address these questions through the application of palaeoecological methods.

### Evidence from the historic (palaeo) environment

Fire has played a central role in shaping landscapes over multi-millennial timescales and has been an important factor in the ecology of UK uplands throughout the Holocene (the last ∼11,700 years) (Davies et al., 2008). Palaeoecological data, such as fossil pollen (a proxy for past vegetation) and microcharcoal (a proxy for past fire activity) from peat sediment cores covering hundreds to thousands of years are widely used tools for reconstructing environmental change (e.g. Innes et al., 2010; Chambers and Daniell, 2011; Garcés-Pastor et al., 2023; Rowney et al., 2023), which reflect local to regional land cover. The global transformation of the biosphere is leading to novel environments, species combinations and ecosystem function (Hobbs et al., 2009) and historical ecology has been used to explore the likelihood of novel ecosystems emerging over time (Woodbridge et al., 2023). We use the term “historic ecological data” to more accessibly represent the term “palaeoecological data” for a wider audience. Such data can provide insights into the trajectories of change in peatlands that led to their current condition, allow exploration of ecosystem resilience and vulnerability to disturbance, and are increasingly applied to conservation issues (Conedera et al., 2009; Davies and Bunting, 2010; Beller et al., 2020).

## Materials and Methods

### Study region

The Peak District National Park (PDNP) (300-400 m above sea level) is an upland protected landscape located in central-northern England (1437 km^2^) that has experienced significant wildfires in recent years (Fig. 1) (Albertson et al., 2009; Barber-Lomax et al., 2022). The area is one of the most southerly and lowest-altitude regions where blanket peat occurs extensively in the UK. Vegetation shifts and human influences on the landscape have been documented in numerous historic (palaeo) ecological studies (e.g. Hicks, 1971; Tallis, 1991; Long et al., 1998; Davies, 2016; Chiverrell, 2001; Simmons et al., 2022) with evidence of forest clearance to provide land for grazing since the Neolithic (Taylor et al., 1994). The PDNP experiences high annual rainfall, is recognised for its biodiversity, and is a significant UK carbon store (Bradfer-Lawrence et al., 2021). Managed burning takes place on red grouse (*Lagopus scotica*) moor, but most wildfires are caused by human actions, either accidentally or on purpose. Wildfires in this region can destroy large areas of habitat, have impacts on wildlife, burn peat, release carbon, impact water and air quality, cause gullying erosion, and affect livelihoods (McMorrow et al., 2009; Barber-Lomax et al., 2022).

**Figure 1.**
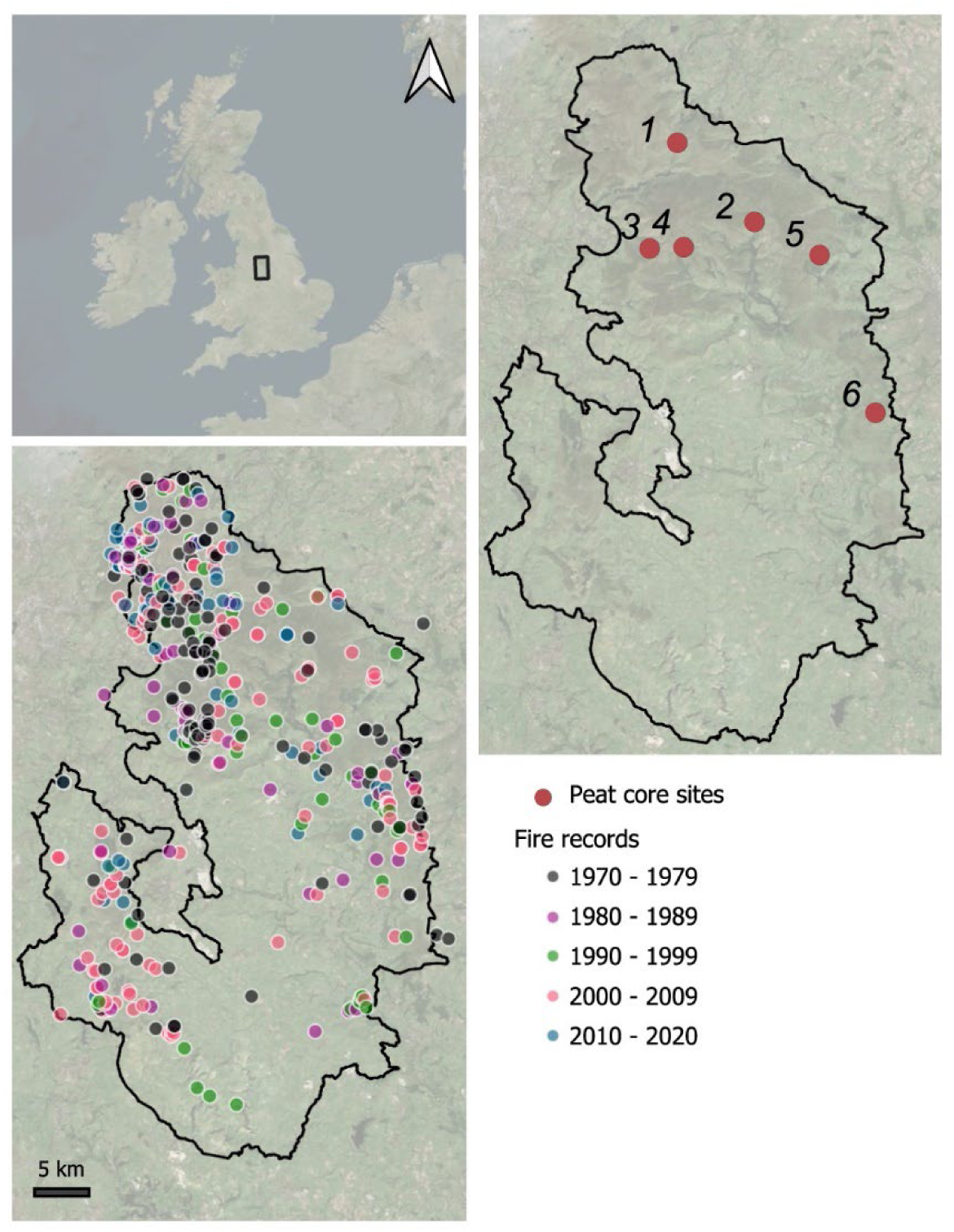
Peak District National Park (PDNP) (black outline). Historic ecological research sites (peat coring locations) illustrated in the right panel (*1: Withens Moor, 2: Cranberry Bed, 3: Coldharbour Moor, 4: Alport Moor, 5: Emlin Dike, 6: Bar Brook*). Fire events (coloured by decade) documented by the Fire Operations Group (PDNP) (left panel).

### Social research methods

To embed local knowledge into the design and interpretation of the historical ecological research, expertise and perspectives were sought from key local stakeholders and actors, such as community members and local organisations. This includes individuals and groups actively involved in making decisions (Ayres et al., 2016) or carrying out activities central to the management and functioning of the environment (see Table 1 for a summary of interviewee roles). Participatory research involves active engagement with individuals who have a stake in the management of the landscape (Davies, 2011; Villamor et al., 2014; Lees et al., 2021). A purposive selection process was used to identify individuals who hold the relevant local knowledge needed for this research (Lees et al., 2021). This approach was used to gather detailed and diverse information from those who are best positioned to provide insights into fire management challenges in the PDNP. This included defining selection criteria (e.g. relevance and depth of experience representing a range of organisations and stakeholder types) and ‘snowball sampling’ to identify participants through networks.

**Table 1.** Interview and workshop participant roles.

| **Workshop** | **Number of participants** |
| --- | --- |
| Local academic | 1 |
| Land agent | 2 |
| Fire and rescue service | 3 |
| National Park authority | 2 |
| Natural resource management organisation | 1 |
| Water supply and wastewater treatment services | 1 |
| **Interviews** |  |
| Moorland management organisation | 1 |
| Multi-agency wildfire management partnership | 1 |
| Fire and rescue service | 5 |
| Public government body | 1 |
| Rural regeneration organisation | 1 |
| Moorland management partnership | 1 |
| Water supply and wastewater treatment services | 2 |
| Land agent | 1 |
| non-profit nature protection charity | 1 |
| *Several individuals also hold roles across multiple organisations including a **multi-agency wildfire management partnership** and **moorland management partnership** |  |

Alongside ongoing conversations with local actors to inform core sampling locations, 14 interviews were conducted with key stakeholders between December 2020 and March 2021. These included land managers, representatives from the PDNP, fire services, water services and other local organisations (Table 1). All interviews were conducted either online or on the telephone and were recorded and transcribed verbatim with confidentiality maintained. An interview guide was used, which aimed to elicit information about understandings of current and past management practices, key challenges to managing wildfire, opportunities for future management, and priorities for different groups. Interview transcripts were analysed thematically following the principles of reflexive thematic analysis (Braun and Clarke, 2022). Codes were generated inductively from the data and integrated into key themes.

### Research questions

All interviewees, along with individuals who were suggested by interviewees, as holding specific knowledge related to the research themes, were invited to a virtual workshop held in May 2021 where preliminary analysis of the interview data and initial analysis of historic ecological data from four of the sites was shared (Fig. 3 a-d and Supplementary Information Table 5). Feedback was gathered and a set of research questions was devised to inform the final analysis and interpretation of the historic ecological datasets: *1) Do fire and vegetation responses vary under different management practices?, 2) How have fire and burning affected landscapes and ecosystems historically in the PDNP?,* and *3) How can long-term historic ecological records inform wildfire management?* In addressing these questions, we use palaeoecological analysis of peat cores to investigate temporal relationships between fire regimes and vegetation (pollen) diversity change, analyse the trends between landscape change, fire activity and peat humification, as an indicator of changing hydrological conditions, and compare trends with a tree ring-inferred precipitation record from the same region.

### Study sites

The research sites were selected to represent priority areas in the PDNP Nature Recovery Plan (PDNPA, 2023) and a range of habitat and management types where fire activity has been concentrated in recent decades (Fig. 1, Table 2 and Supplementary Information 1). This study includes two newly analysed sites (Coldharbour Moor and Alport Moor) and four previously published (Davies, 2016) sites (Emlin Dike, Withens Moor, Bar Brook and Cranberry Bed). Coldharbour Moor and Alport Moor are Special Areas of Conservation (SACs) representing degraded but recovering peatland habitats, characterised by heather-dominated moorland and blanket bog, with ongoing restoration through re-vegetation and re-wetting of degraded areas (National Trust, 2018). Re-vegetating of degraded moorlands involves species such as *Sphagnum* mosses, cotton grasses (*Eriophorum spp*.), and heathers (*Calluna vulgaris*), which are essential for rebuilding peat structure and promoting long-term ecosystem recovery (Lunt et al., 2010). Emlin Dike and Withens Moor, both within the Dark Peak SSSI, include fen and blanket peat habitats. Emlin Dike has been managed through rotational burning for grouse production since AD 1950 (Davies, 2016). In the region there have been conservation actions to restore native vegetation and control purple moor grass (*Molinia caerulea*) (Natural England, 2009). Purple moor grass is considered a major threat to moorland conservation in the UK and elsewhere in Europe (Marrs et al., 2004; Pilkington et al., 2021). Cranberry Bed and Bar Brook, also part of the Dark Peak SSSI, comprise fen and channel mire habitats on deep peat with varied hydrological conditions (Natural England, 2009; Davies, 2016).

**Table 2.** Peat coring locations in the Peak District National Park. Site numbers 1 to 6 correspond with numbers shown on the map in Figure 1.

| Site name | Elevation (m a.s.l.) | Latitude | Longitude | Reference | Site characteristics |
| --- | --- | --- | --- | --- | --- |
| Coldharbour Moor (3) | 457 m | 53.43391 | -1.88548 | This study | Special Area of Conservation (SAC) characterised by extensive heather-dominated peat moorland showing visible evidence of recent fire events. The site is considered degraded due to historic overgrazing, burning, and industrial pollution, though ongoing conservation work aims to promote recovery through re-vegetation and re-wetting initiatives. |
| Alport Moor (4) | 475 m | 53.43488 | -1.83951 | This study | SAC within the North Peak Environmentally Sensitive Area (ESA) scheme, encompassing blanket bog, dry heathland, and mires interspersed with woodland patches. Vegetation is dominated by heather (*Calluna vulgaris*), cotton grasses (*Eriophorum spp.*), and *Sphagnum* mosses, with restoration efforts focused on improving peat structure and enhancing hydrological conditions (re-vegetation and re-wetting). |
| Emlin Dike (5) | 390 m | 53.42861 | -1.65583 | Davies (2016) | Part of the Dark Peak Site of Special Scientific Interest (SSSI). Consists of a fen with areas of deeper peat located on heather-rich moorland. The site holds favourable conservation status and has been managed through rotational patch burning for grouse production since the mid-twentieth century. |
| Withens Moor (1) | 440 m | 53.51889 | -1.84806 | Davies (2016) | The site is within the Dark Peak SSSI and comprises deep blanket peat and acid grassland dominated by purple moor grass (*Molinia caerulea*). The site has 'unfavourable recovering' conservation status, with management efforts targeting the control of *Molinia* and the reintroduction of native species, including *Sphagnum* mosses. |
| Cranberry Bed (2) | 290 m | 53.45528 | -1.74389 | Davies (2016) | Fen located at the base of a valley slope within the Dark Peak SSSI. A habitat previously classified as having 'unfavourable recovering' conservation status. The site supports a mix of peat-forming vegetation and reflects the influence of hydrological variation. |
| Bar Brook (6) | 360 m | 53.30139 | -1.58028 | Davies (2016) | A channel mire situated on intact deep peat within the Dark Peak SSSI. The site's vegetation includes wet and dry heath, purple moor grass, and associated grassland, representing a mosaic of peatland habitats with ongoing ecological interest. |

### Historic (palaeo) ecological methods

Peat cores were collected using a Russian corer (Table 2) and sub-sampled at high-resolution for fossil pollen, micro-charcoal and peat humification analysis. Standard methods for pollen analysis and identification (Andrew, 1984; Moore et al., 1991) were applied to one cm^3^ peat sub-samples. A *Lycopodium* marker tablet was added to each sample for calculation of pollen and charcoal concentrations (Stockmarr, 1971). Samples were heated in 10% potassium hydroxide (KOH) to remove humic acid, sieved to retain the 10-180 mm fraction, acetolysis digestion was applied to remove non-pollen organics, and samples were mounted in silicon oil. A minimum of 300 terrestrial pollen grains were identified under a light microscope and counted for each sample alongside aquatics and spores. Charcoal fragments were recorded and counted within pollen samples and separated in two size classes (<50 µm and >50 µm) to separate local and regional fire signals (Tinner and Hu 2003). One cm^3^ peat samples were taken for radiocarbon (^14^C) dating and analyses were conducted by either the Chrono Centre (Queen’s University Belfast) or NERC’s National Environmental Isotope Facility. In this study, peat humification was analysed for the cores from Alport Moor and Coldharbour Moor by measuring the percentage of light that is transmitted through a solution of the peat using a spectrophotometer (Blackford and Chambers, 1993). Lower absorbance values can indicate drier climatic conditions while higher absorbance values indicate wetter climatic conditions (Charman, 2002). A tree ring-inferred precipitation record (oxygen isotope ratios) reflecting May-August rainfall for England and Wales, which shows good agreement with instrumental rainfall records (Loader et al., 2020), is compared with the historic ecological datasets to explore possible relationships between climate trends (precipitation), vegetation and fire activity.

Peat core chronologies were constructed using the ‘rbacon’ (v. 3.5.2) (Blaauw et al., 2021) Bayesian chronological modelling R package. The four cores from Davies (2016) were dated using AMS radiocarbon ages, lead-210 and spheroidal carbonaceous particles (SCPs) (soot particles from fossil fuel combustion) (Davies, 2016) to construct chronologies. Age-depth models (Fig. 2 a-b) were produced for Coldharbour Moor and Alport Moor using 14 radiocarbon dates (Table 3, Fig. 2) to produce chronologies, which were converted to calendar years. The core surface age was set as −71 yr BP ±30. Other age-depth modelling settings included mean accumulation rate to guide expected sedimentation, memory parameters to control the degree of autocorrelation in accumulation between adjacent sections, and a mean hiatus length to allow for the possibility of brief depositional gaps (Blaauw & Christen, 2011). The age-depth models for Cranberry Bed, Emlin Dike, Withens Moor and Bar Brook are described and presented in Davies (2016). Individual taxa are presented as a percentage of the total terrestrial pollen assemblage. Charcoal concentration (fragments/cm^2^) was calculated using the ratio of *Lycopodium* spores added to the sample, and charcoal influx (fragments/cm^2^/yr) was calculated by dividing concentration by the sediment accumulation rate according to the age-depth models. Charcoal data are presented as ‘influx’ (i.e. deposited charcoal fragments per unit area and time) because this approach allows for comparison between sites and accounts for variation in sediment accumulation rates (Mooney & Tinner, 2011). The charcoal fractions above and below 50 µm were combined for calculation of charcoal influx to represent a collective local and regional fire signal.

**Figure 2.**
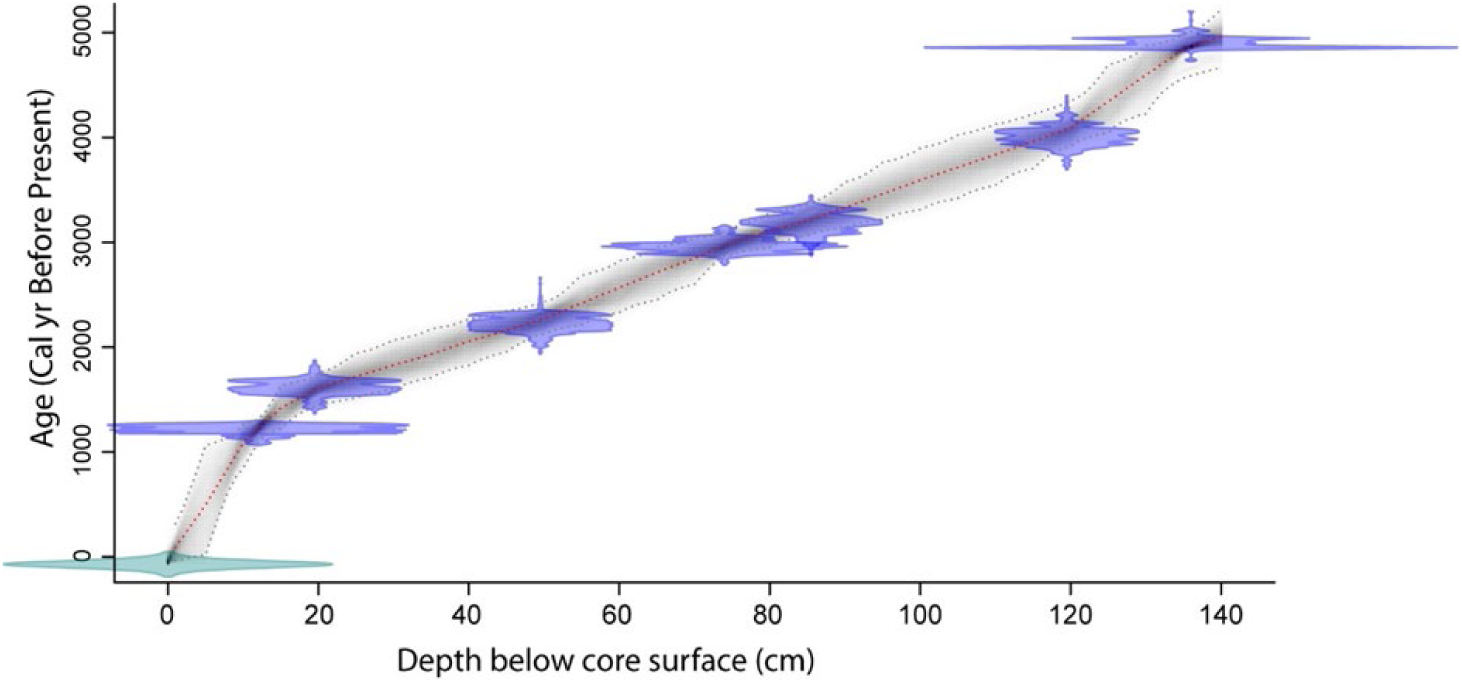
**a)** Age-depth model for Coldharbour Moor based on seven radiocarbon (^14^C) dates. Shaded areas represent error within the ^14^C dating (blue) and age-depth modelling (grey).

**Figure 2.**
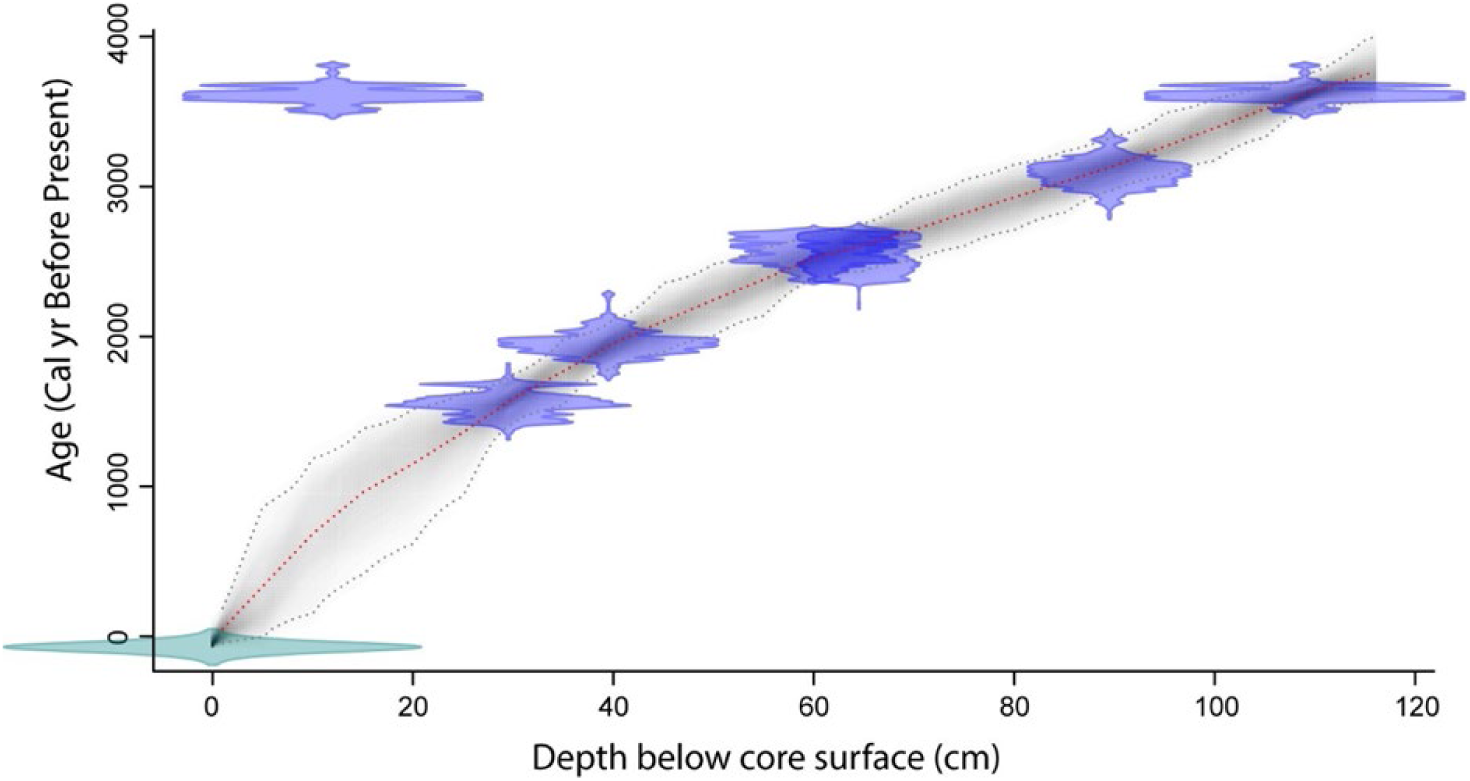
**b)** Age-depth model for Alport Moor based on seven radiocarbon (^14^C) dates. Shaded areas represent error within the ^14^C dating (blue) and age-depth modelling (grey).

**Table 3.**
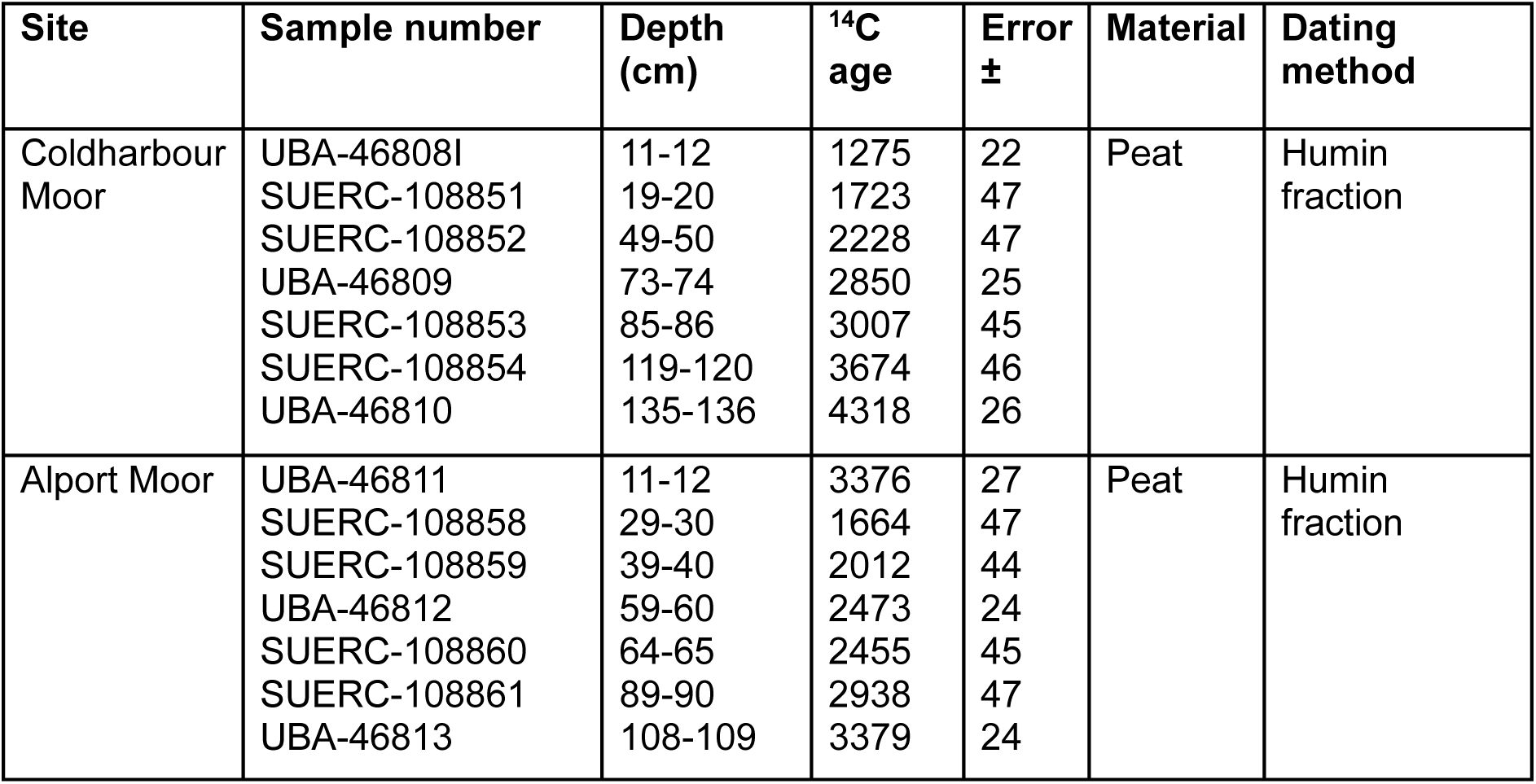
Radiocarbon (^14^C) dates for peat sediment cores from Alport Moor and Coldharbour Moor.

| Site | Sample number | Depth (cm) | $^{14}\text{C}$ age | Error ± | Material | Dating method |
| --- | --- | --- | --- | --- | --- | --- |
| Coldharbour Moor | UBA-46808I | 11-12 | 1275 | 22 | Peat | Humin fraction |
|  | SUERC-108851 | 19-20 | 1723 | 47 |  |  |
|  | SUERC-108852 | 49-50 | 2228 | 47 |  |  |
|  | UBA-46809 | 73-74 | 2850 | 25 |  |  |
|  | SUERC-108853 | 85-86 | 3007 | 45 |  |  |
|  | SUERC-108854 | 119-120 | 3674 | 46 |  |  |
|  | UBA-46810 | 135-136 | 4318 | 26 |  |  |
| Alport Moor | UBA-46811 | 11-12 | 3376 | 27 | Peat | Humin fraction |
|  | SUERC-108858 | 29-30 | 1664 | 47 |  |  |
|  | SUERC-108859 | 39-40 | 2012 | 44 |  |  |
|  | UBA-46812 | 59-60 | 2473 | 24 |  |  |
|  | SUERC-108860 | 64-65 | 2455 | 45 |  |  |
|  | SUERC-108861 | 89-90 | 2938 | 47 |  |  |
|  | UBA-46813 | 108-109 | 3379 | 24 |  |  |

Pollen diversity measures represent both taxa richness and how evenly different types of plants are represented in the sample (Birks et al., 2016). A Shannon diversity index score was calculated for each pollen sample using untransformed pollen percentage data for all terrestrial pollen taxa via the R package ‘vegan’ (Oksanen et al., 2019). The index captures vegetation heterogeneity providing a measure of pollen-type diversity, which reflects landscape diversity (Matthias et al., 2015; Woodbridge et al., 2021). Higher values represent diverse and equally distributed communities while lower values represent less diverse and less equally distributed vegetation. For a description of the process involved in producing Shannon index scores from pollen data see Woodbridge et al. (2021). Spearman’s rank correlation coefficient was used to identify relationships between the datasets. Pollen samples were analysed using Canonical Correspondence Analysis (CCA) and Principal Components Analysis (PCA) using the R package ‘vegan’ (Oksanen et al., 2019) and ordination plots were used to explore relationships with other variables (charcoal, Shannon diversity and SCPs). PCA plots for the four previously published sites are presented in Davies (2016) supplementary materials. An analogue-matching approach was also applied to the pollen datasets to identify similar assemblages between the modern (or most recent) pollen sample and all other samples in the same dataset using the R package ‘analog’ (Simpson and Oksanen, 2021). Analyses were applied to all land pollen taxa in the datasets (see Supplementary Information 2 a-e for pollen diagrams showing all land taxa and *Sphagnum* for each site). A selection of taxa occurring at above 5% abundance are presented in Fig. 3.

## Results

### Social-cultural knowledge

Through participatory research, several commonly emerging themes were identified surrounding the impacts of wildfire, management practices, fire forecasting, and fire-hazard awareness (Supplementary Information Table 6). These interacting issues can be categorised into environmental, social and cultural, and management issues. Many participants discussed the environmental impacts of wildfires, which includes loss of biodiversity, localised extinctions of species, displacement of wildlife, and resulting pollution.

Often, environmental impacts are linked to social issues, for example, resulting water pollution must be managed by water companies after fires to ensure that drinking water remains safe. Furthermore, wildfire events are often clustered near to public open access areas, demonstrating the importance of understanding and incorporating public values into planning. Interviewees reflected that progress is needed in raising awareness among the public; they identified the need for more education around the risk of wildfire and consistent messaging in visitor hotspots. In the PDNP there are collaborative initiatives for wildfire management in place, such as the Moors for the Future Partnership and the PDNP Fire Operations Group (FOG), which comprises landowners, gamekeepers, fire services, water companies, and conservation bodies coordinating wildfire response and prevention. Many interviewees valued these initiatives and the relationships they foster. Economic challenges were also identified, particularly the impact on tourist and leisure activities, costs required for restoration following wildfires, and immediate costs to manage specific fire events.

Challenges were identified around issues of funding and resources for fire mitigation and recovery, alongside the periodicity of fires, for example, less frequent fires lead to decreased resources for wildfire management, which then leads to fuel build-up, and larger fires consequently. The need for strategic contextualised approaches to fire management and risk, appropriate land management, recognition of the multiple uses of the land, recognition of land managers’ roles, identification of the causes of fire, and the importance of appropriate local-scale policy and risk mitigation were highlighted:

> “*We have a unique habitat and unique land use, multiple land use, you know, we’ve got grazers, landowners, sporting rights, mineral rights, they’re all layered on top of each other all on one patch of land. So it’s highly complex, and unique*.” (Interviewee)

> “*Quite often we’ll have a policy that comes down the line from government and unless it’s met with people’s experience from the ground up, quite often misses the mark, and you get unintended consequences.”* (Interviewee)

Participatory research revealed that there is a great deal of scientific research taking place in the PDNP, but this needs to be more frequently embedded into policy and guidance, which in turn would benefit from greater input from those who manage the land:

> *“Trying to put together the practitioner with the scientist is really important, and the scientists must have, you know, due regard, respect for the view of the practitioner because otherwise, you know, we’re in for- you only need to look at things like the Staybridge fire to realise something is amiss.”* (Interviewee)

When interviewees were asked about knowledge gaps in current understanding of environmental responses to fire, topics that arose included the need to understand how fire and burning have affected landscapes and ecosystems, the need to recognise patterns of regeneration following fire, and whether these responses vary under different management practices. The most commonly occurring words used by interviewees are summarised in Table 4b. Words such as “people”, “management”, “volunteers”, “partnership” and “coordinated” illustrate the role of communities and stakeholders in fire management, and “peat”, “woodland” and “nesting” indicate concerns over the impacts of fires, while “work”, “resources” and “equipment” demonstrate the inputs that are required for management, and words such as “gatekeepers” and “fuel” reflect some of the barriers to effective management. The main causes of fire ignition in the PDNP involve human action (e.g. controlled burns to manage fuel loads and vegetation). Ignition sources listed in the FOG fire monitoring data for fires recorded in recent decades include accident/arson, recreational campfires, cigarettes, contractor operations, controlled burns, forestry fires, parachute flare (military training), and uncontrolled burns. Furthermore, environmental conditions play an important role in ignition risk and fire spread (e.g. heatwaves, drought and wind).

**Table 4a.**
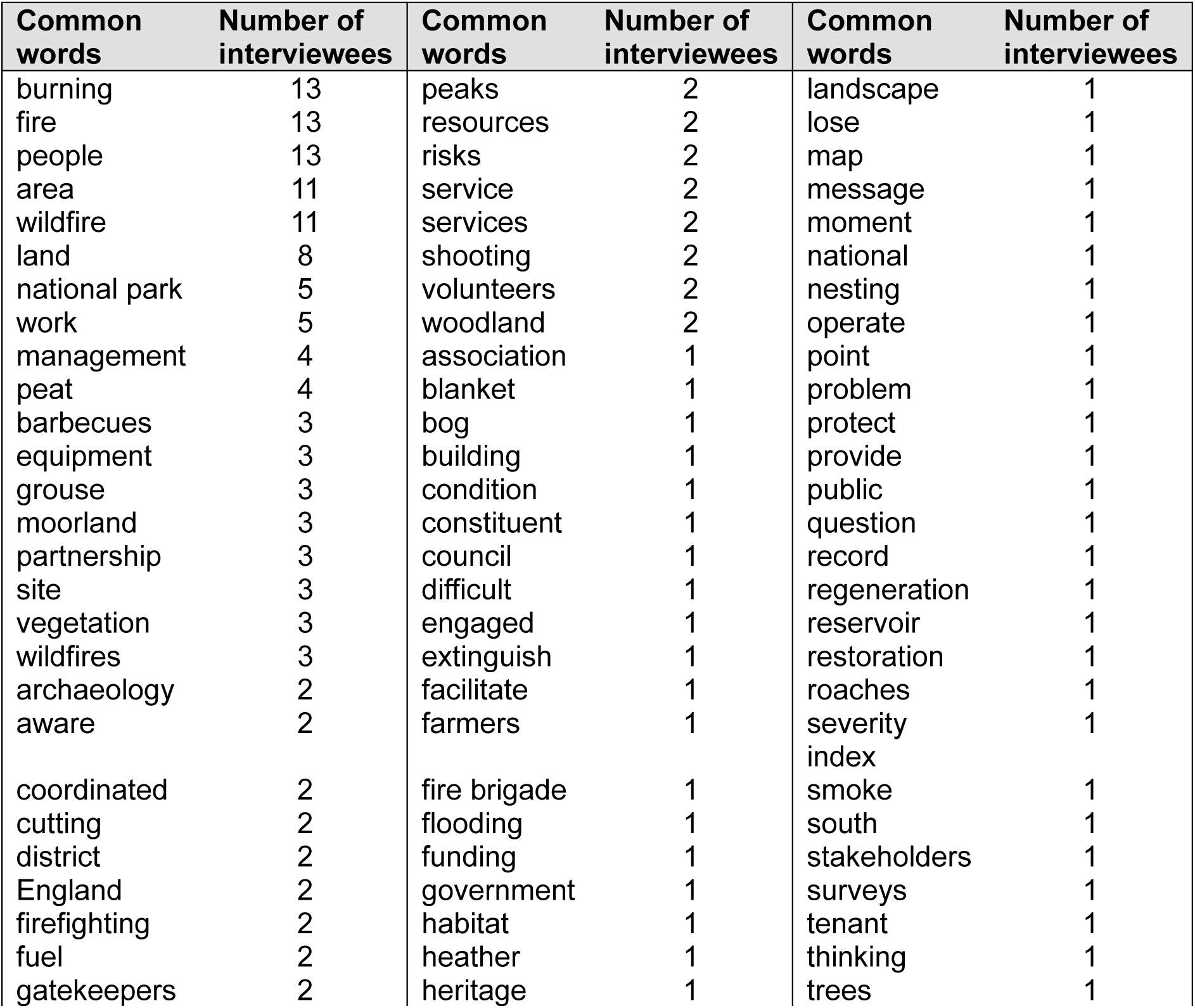
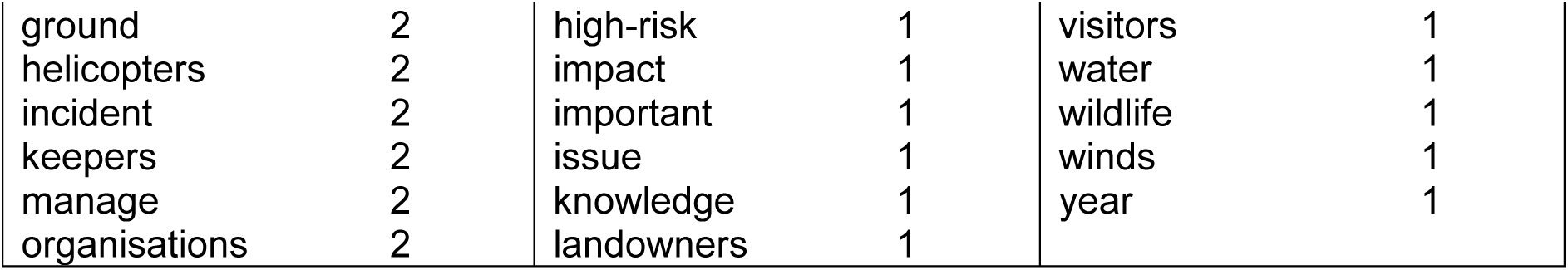
Commonly occurring words identified in interview transcripts with the number of interviewees that the words were commonly used by.

**Table 4b.** Commonly occurring themes identified in interview transcripts with the number of interviews that the themes arose within.

| Theme | Number of Interviewees |
| --- | --- |
| Wildfire impacts | 14 |
| Climate and weather | 14 |
| Land use conflicts | 14 |
| Monitoring and preparedness | 14 |
| Historical knowledge and management | 14 |
| Fuel load and fire risk | 13 |
| Policy and regulation | 13 |
| Peatland rewetting and restoration | 12 |
| Vegetation and biodiversity | 12 |
| Prescribed burning | 11 |

### Historic ecological change

Although the spatial extent of the fires in relation to the peat bogs was not explicitly investigated in the past, the presence of charred remains of local vegetation during field sampling suggests recent burning activity that may have affected both the surrounding areas, and the peat surfaces themselves. Comparison of wildfire records obtained from the PDNP local FOG for decadal time-windows (1970-2020) revealed dissimilar patterns between these periods (Fig. 1) and indicate that fires have been clustered in the northern, western and eastern areas of the PDNP for the last 70 years. During the decade from 2010 to 2020, most fires were concentrated in the northern reaches of the PDNP with sporadic fires persisting in other areas. The pollen records (Fig. 3 a-f) indicate continual abundant cover of heath and grasses and various low abundance herbaceous types with presence of peat moss (*Sphagnum*) and sedges (*Cyperaceae*) alongside deciduous trees and pine plantations in the 19^th^ - 20^th^ century. *Sphagnum* spores are produced in lower abundance than the pollen of vascular plants, therefore values in the diagrams do not reflect land cover. Pollen percentage data reflect vegetation growing within a few kilometres of a coring site, but the pollen signal will be dominated by plants growing on and immediately adjacent to the sampled peatland (Jacobson and Bradshaw, 1981).

**Figure 3.**
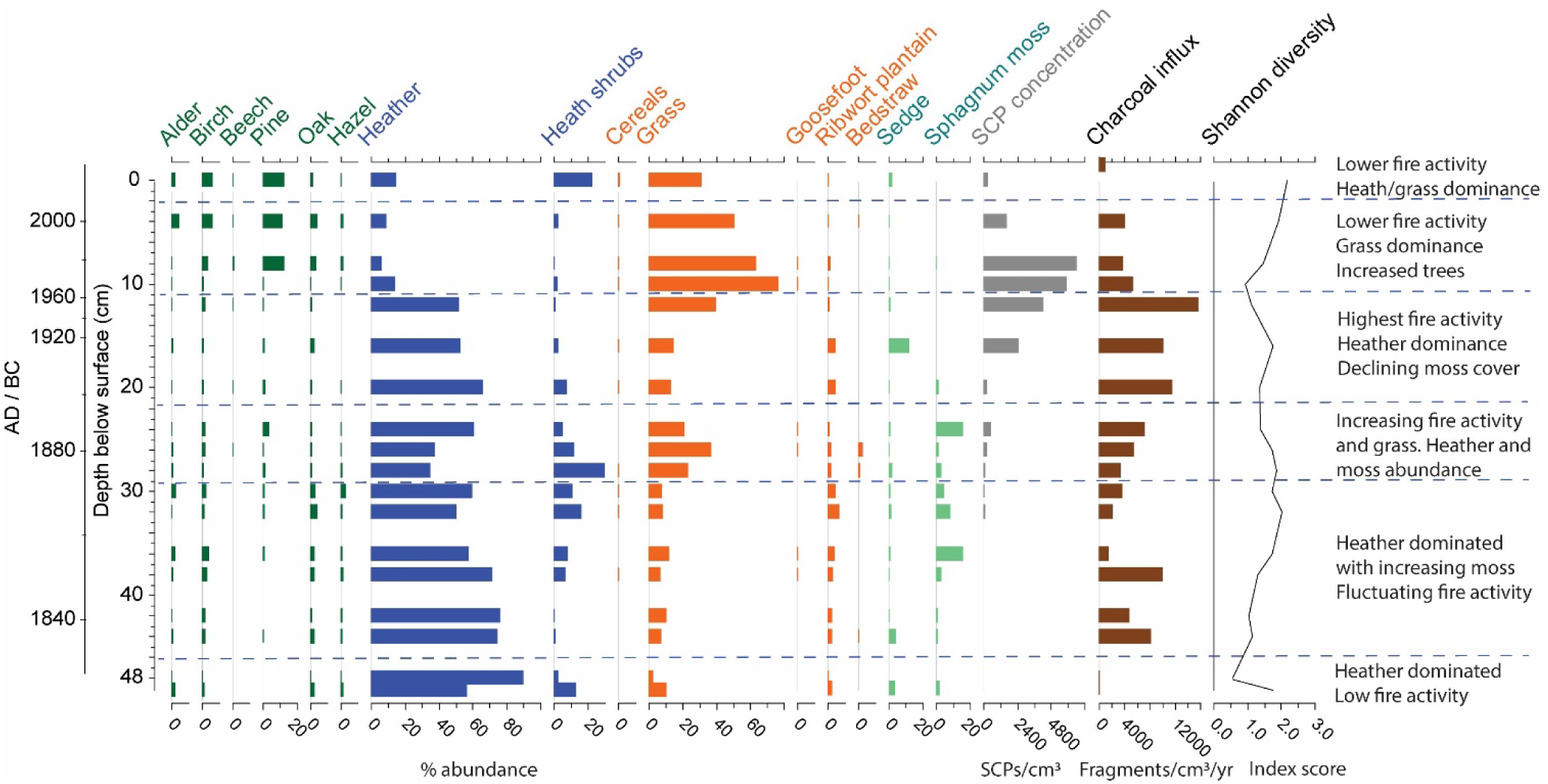
**a)** Cranberry Bed percentage data for selected pollen taxa with spheroidal carbonaceous particles (SCP) concentration, charcoal influx (fragments/cm^3^/yr) and pollen-inferred Shannon diversity index. Green bars: trees, blue bars: heathland, orange: herbs, sage: moss and sedge, and brown: charcoal.

**Figure 3.**
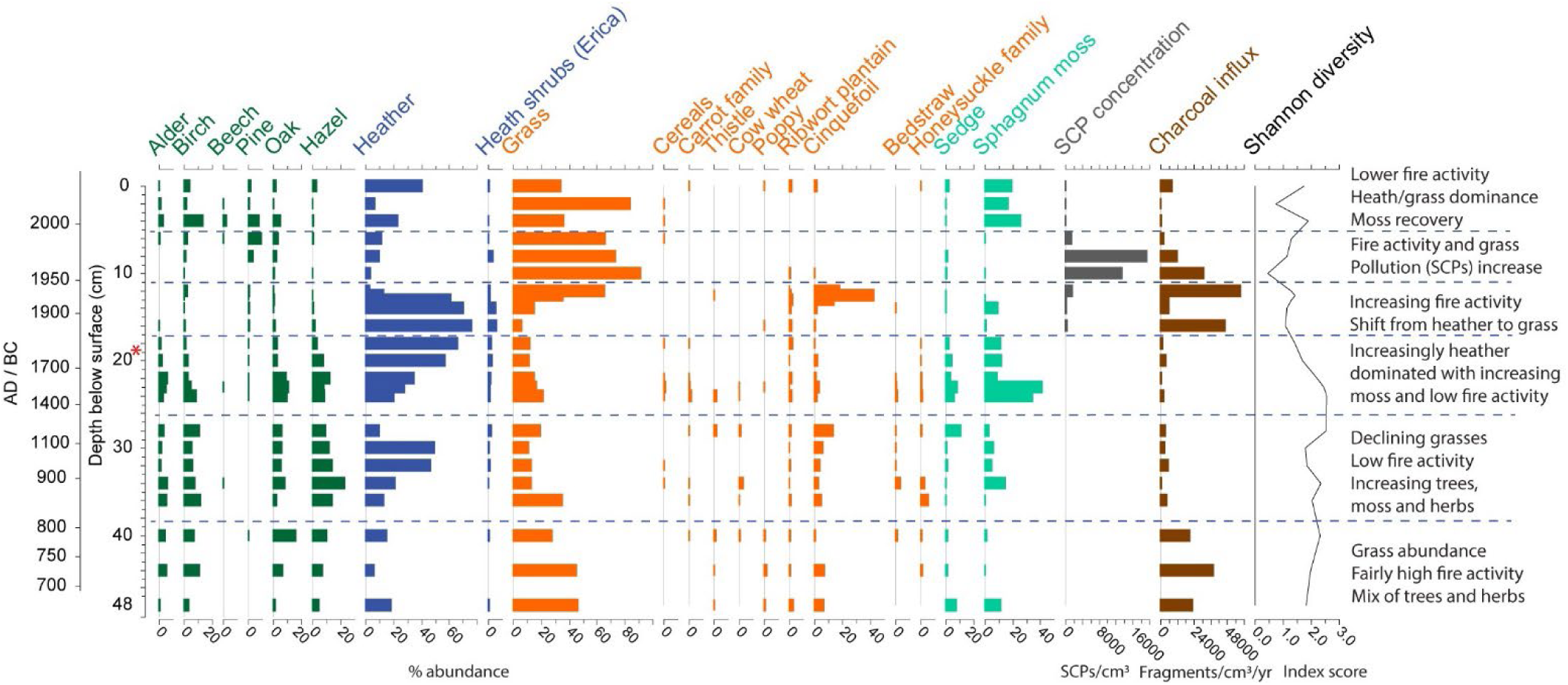
**b)** Bar Brook percentage data for selected pollen taxa with spheroidal carbonaceous particles (SCP) concentration, charcoal influx (fragments/cm^3^/yr) and pollen-inferred Shannon diversity index. Red asterisk highlights modern analogue sample. Green bars: trees, blue bars: heathland, orange: herbs, sage: moss and sedge, and brown: charcoal.

**Figure 3.**
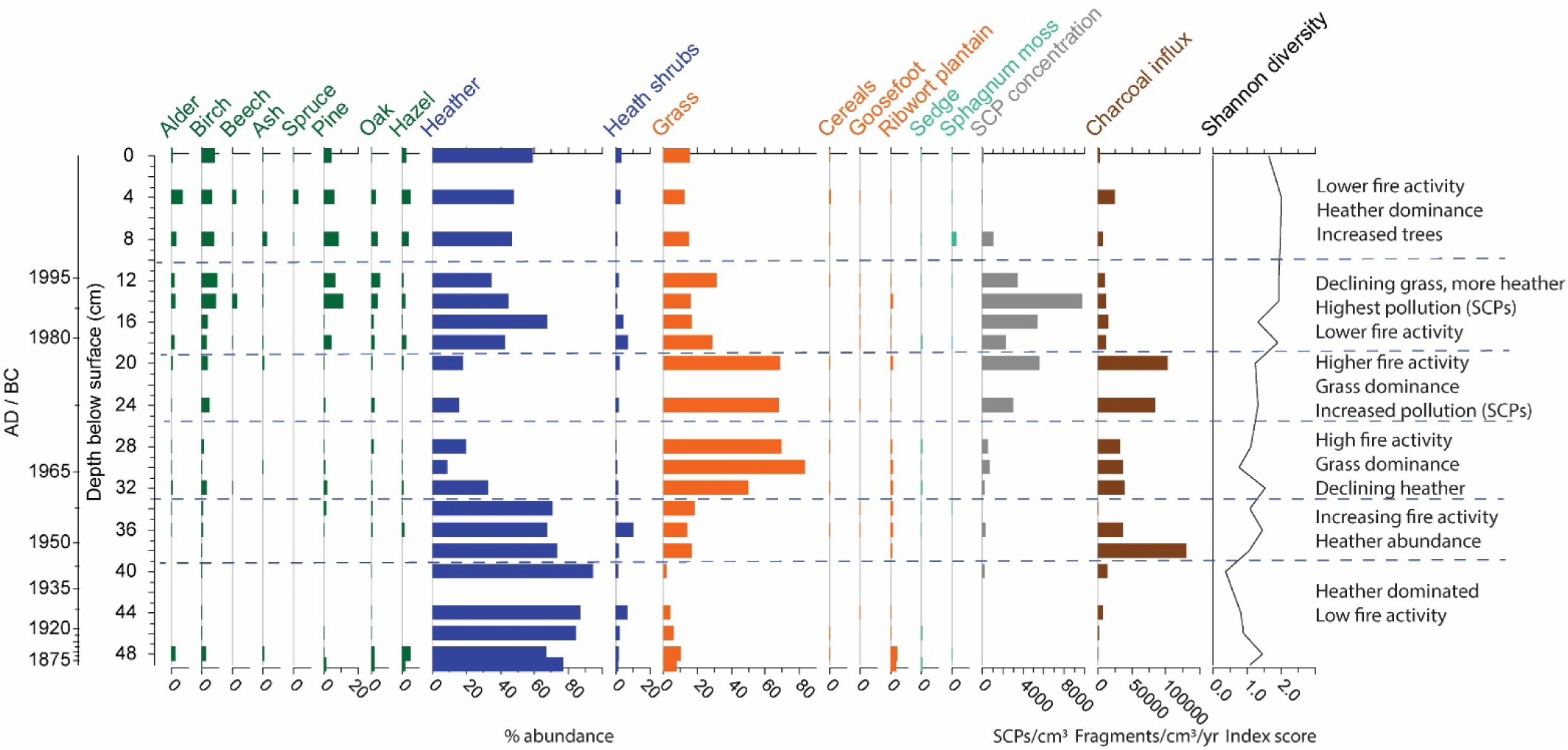
**c)** Emlin Dike percentage data for selected pollen taxa with spheroidal carbonaceous particles (SCP) concentration, charcoal influx (fragments/cm^3^/yr) and pollen-inferred Shannon diversity index. Green bars: trees, blue bars: heathland, orange: herbs, sage: moss and sedge, and brown: charcoal.

**Figure 3.**
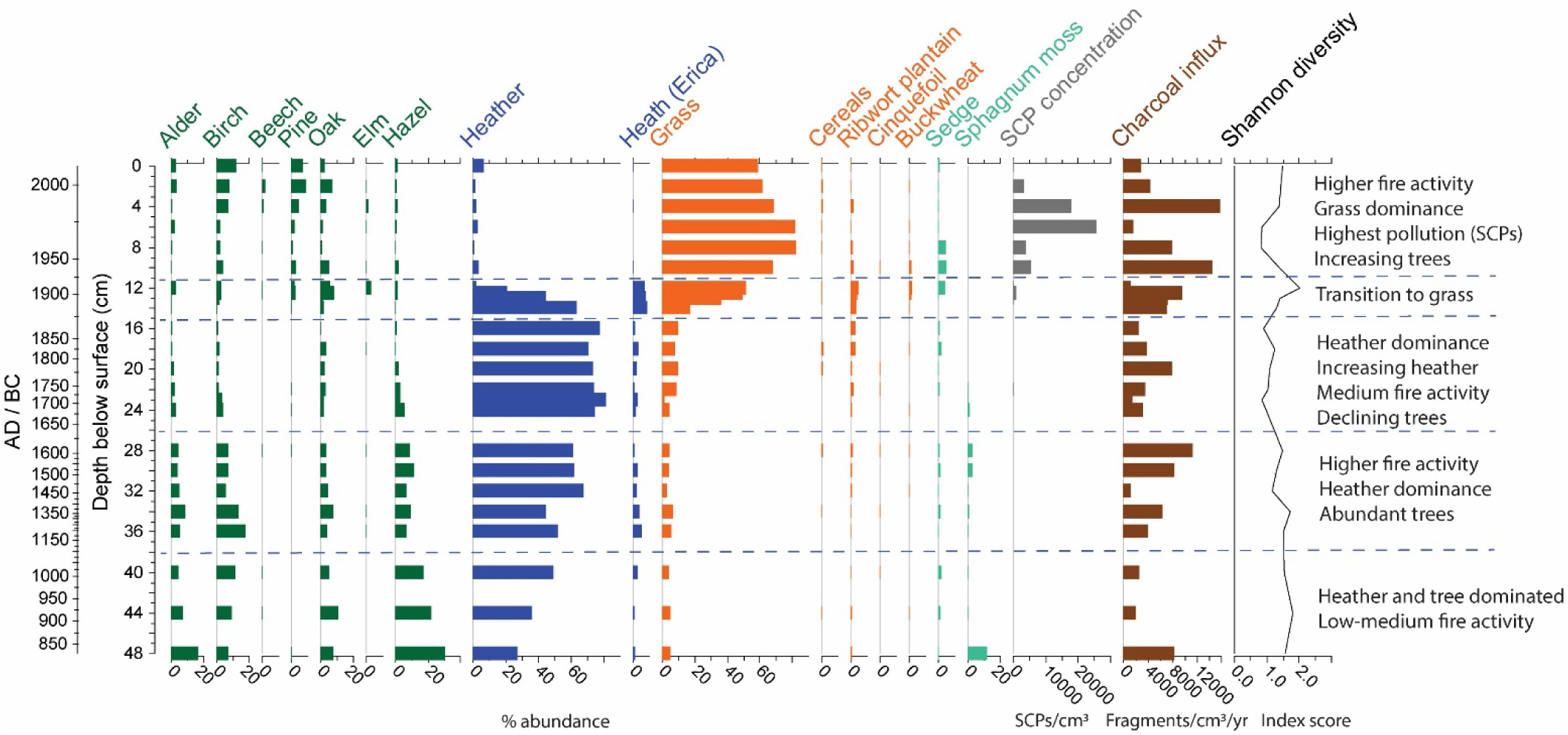
**d)** Withens Moor percentage data for selected pollen taxa with spheroidal carbonaceous particles (SCP) concentration, charcoal influx (fragments/cm^3^/yr) and pollen-inferred Shannon diversity index. Green bars: trees, blue bars: heathland, orange: herbs, sage: moss and sedge, and brown: charcoal.

**Figure 3.**
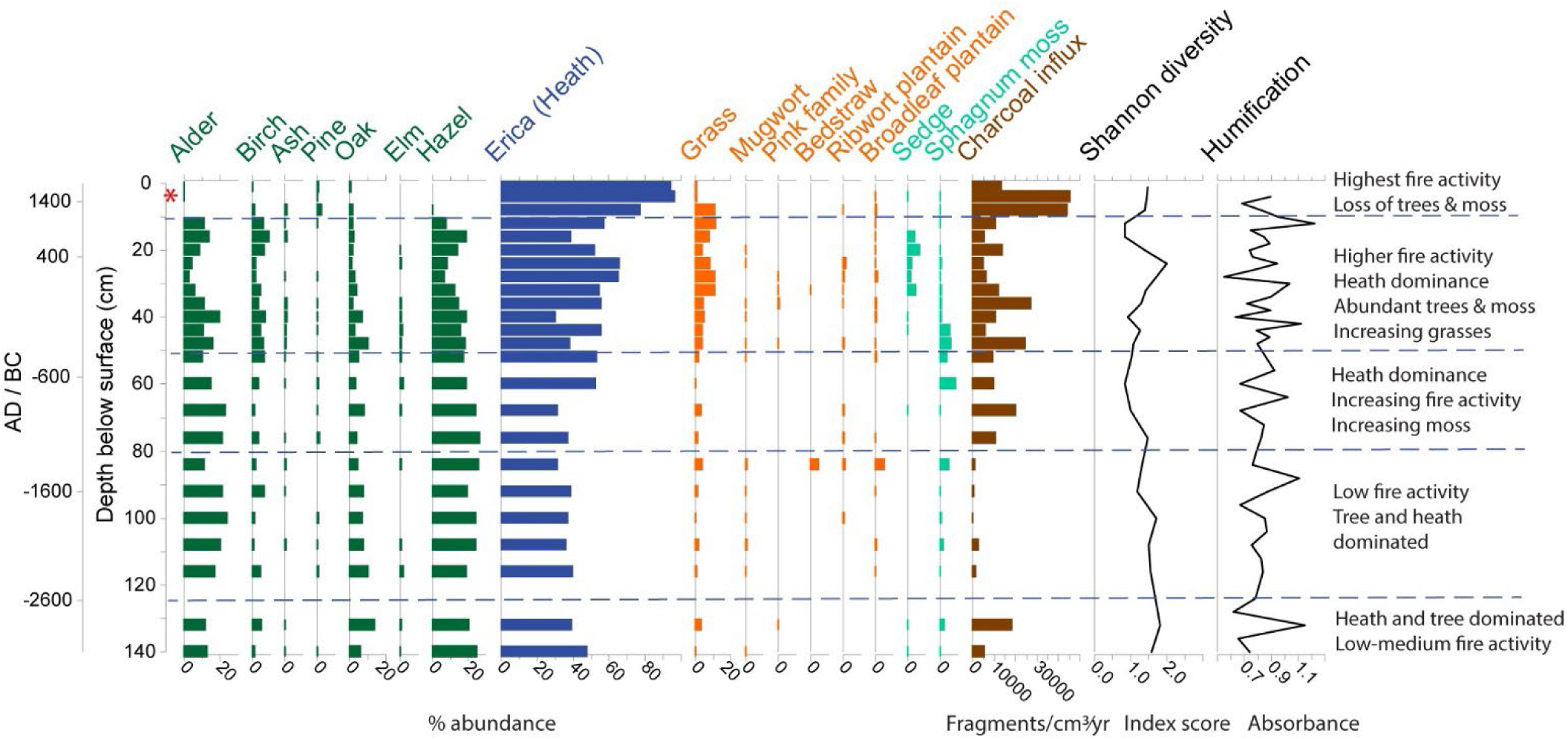
**e)** Coldharbour Moor percentage data for selected pollen taxa with charcoal influx (fragments/cm^3^/yr), pollen-inferred Shannon diversity index, and peat humification (detrended absorbance values). Red asterisk highlights modern analogue sample. Green bars: trees, blue bars: heathland, orange: herbs, sage: moss and sedge, and brown: charcoal.

**Figure 3.**
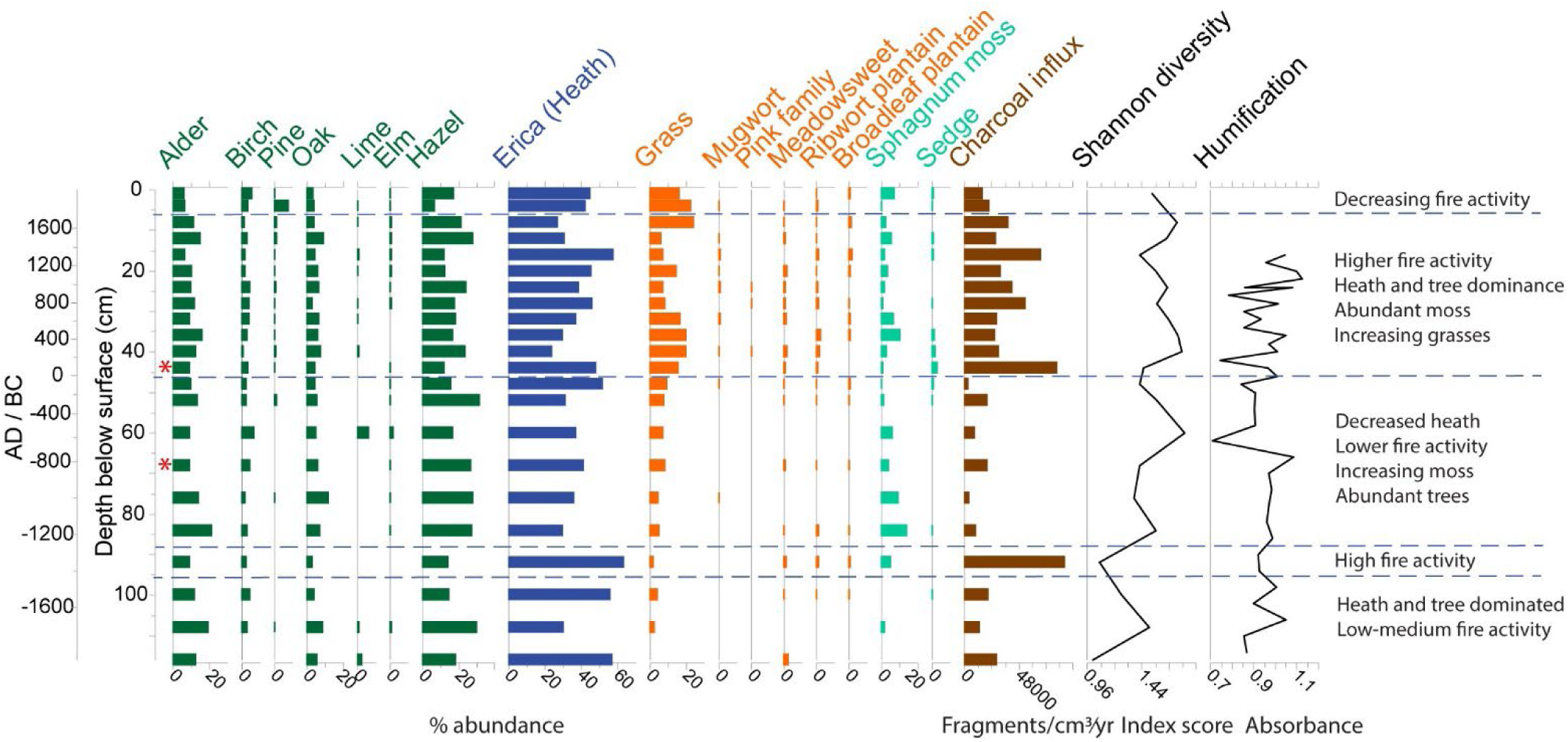
**f)** Alport Moor percentage data for selected pollen taxa with charcoal influx (fragments/cm^3^/yr), pollen-inferred Shannon diversity index, and peat humification (detrended absorbance values). Red asterisks highlight modern analogue samples. Green bars: trees, blue bars: heathland, orange: herbs, sage: moss and sedge, and brown: charcoal.

The six historic ecological records capture legacies of past environmental change and human land use and illustrate that fire has been a persistent feature of the landscape through time (Fig. 3 a-f). In the record from Cranberry Bed (Fig. 3a), which covers the most recent ∼200 years, fire activity persists through the entire record showing a gradual increase. At this site, a fire tolerance threshold appears to have been reached as the system switches from heather to grass dominance (∼AD 1960) and a recent (late 20^th^ century) decline in fire activity is accompanied by recovery of heathland species and tree/shrub cover. Fire activity was generally lower in the earlier record from Bar Brook (Fig. 3b) (dating from ∼AD 750) with increasing fire activity from the late 19^th^ century. *Sphagnum* moss is now abundant at this site indicating favourable ecological conditions for peat formation. Similarly, Milner et al. (2021) identified tipping points in a UK peatland system where a degraded state was followed by a return to peat forming conditions and restored ecological functions. The record from Emlin Dike (Fig. 3c) (dating from ∼AD 1875), shows increasing fire in the 20^th^ century similarly associated with loss of heather and increasing grass cover, which is followed by decreasing fire activity and recovery of heather.

Withens Moor (Fig. 3d) (dating from ∼AD 850) has experienced higher and constant fire activity, with peaks in the 20^th^ century, and significant loss of heather and persistent grass dominance since ∼AD 1920. The record from Coldharbour Moor (Fig. 3e) covering an earlier period (from ∼3000 BC) captures the time when heathland and tree cover was significantly higher with low grass cover. As fire activity increased, grasses became more abundant, but this level of fire activity was not significant enough to cause vegetation assemblage changes as captured in the records covering recent centuries. At Alport Moor (Fig. 3f) (dating from ∼1800 BC), an increase in fire activity is reflected by grass dominance, but not at the values shown in the four more recent records. Although the records indicate a 20^th^ century peak in fire activity, the different timings and intensities of changing fire regimes across sites highlight that fire is regionally diverse and influenced by site level characteristics. The tree-ring inferred precipitation reconstruction (Loader et al., 2020) shows a gradually decreasing trend since ∼AD 1750, which corresponds with increasing fire activity in the charcoal records (Fig. 4). Changing fire activity also relates to shifting peat and vegetation moisture levels (Davies et al., 2023). A drier period indicated by the humification data may have preceded the trend towards increased fire activity.

**Figure 4.**
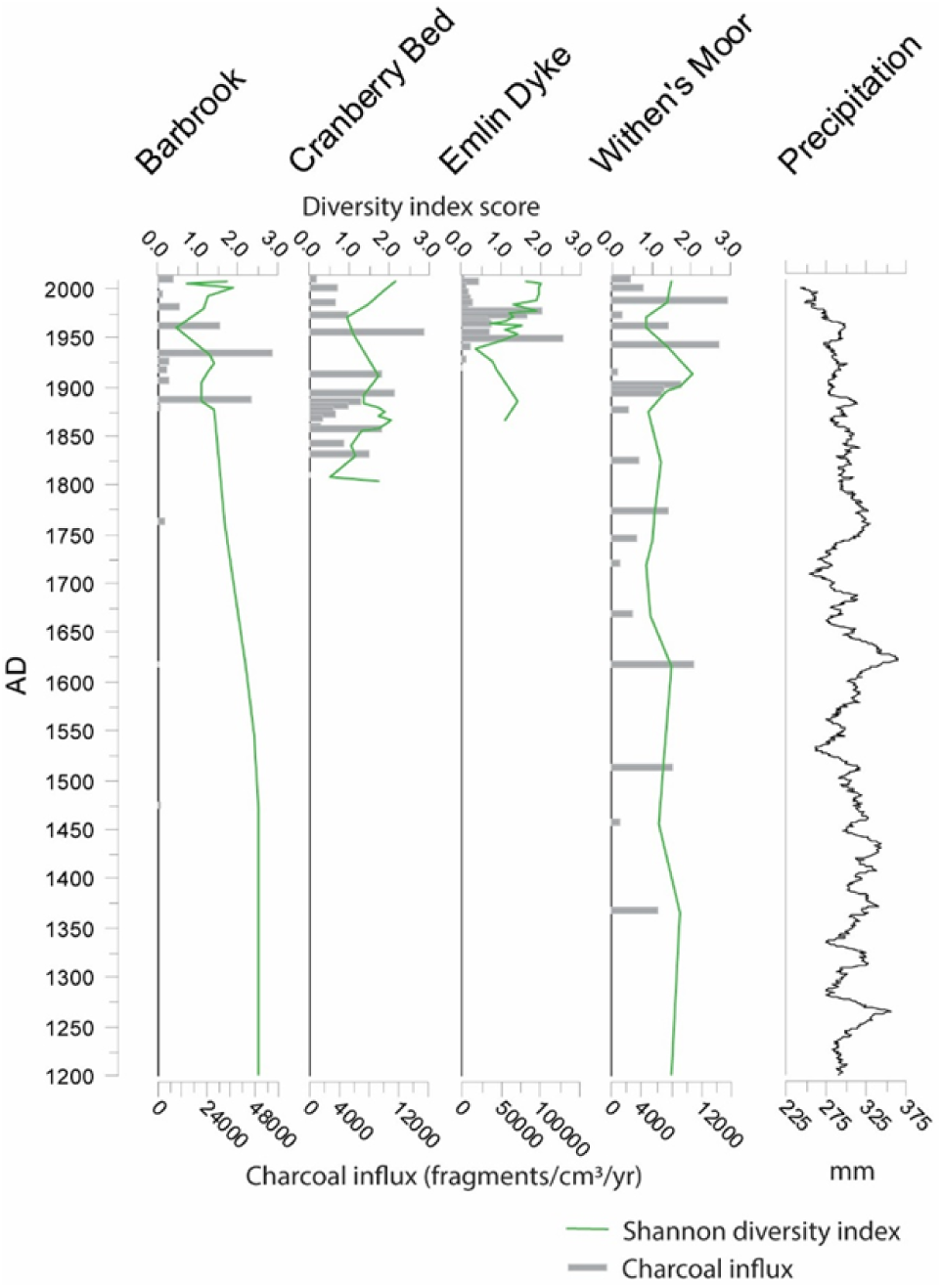
Charcoal influx (fragments/cm^3^/yr) and pollen-inferred Shannon diversity index with tree ring inferred precipitation (May-August) (Loader et al., 2020).

In addition to naturally occurring fire, burning has been used as a tool to control vegetation over long time periods. Since ∼1300 BC in the record from Alport Moor this is indicated by co-occurrence of grazing indicators (ribwort plantain: *Plantago lanceolata*) and increased charcoal. At Coldharbour Moor, burning may have been used as a method for vegetation control since ∼1000 BC, as indicated by grazing pollen indicators and increased fire activity. Numerous statistically significant relationships have been identified between pollen assemblage change, charcoal influx, Shannon (pollen) diversity, SCP concentration (as an indicator of pollution) and peat humification (as a hydrological proxy) (Supplementary Information Table 5). The records from Alport Moor and Coldharbour Moor signify that during the latter mid-Holocene (∼3000 BC onwards) fire led to promotion of herbaceous plant types, but the more recent records indicate that fires are now having negative impacts on the diversity of herbaceous plants (since ∼AD 1950). At Bar Brook there is a negative relationship between charcoal and *Sphagnum* moss, cereal crops, and other herbs. Similarly, there is a negative relationship between charcoal influx and *Sphagnum* moss at Emlin Dike, implying that fire increase may be associated with peat moss decrease. However, research has also shown loss of *Sphagnum* species to be linked to intensification of land-use (drainage, peat cutting, grazing, erosion) and climate change (Hughes et al., 2008).

At Cranberry Bed, a negative relationship exists between charcoal influx and vegetation evenness implying that fire activity has led to more homogeneous vegetation. While correlation does not necessarily indicate causation and responses likely relate to multiple factors, these results suggest that the change in fire regimes between the latter mid-Holocene and recent centuries is an important factor in moorland vegetation dynamics. An analogue matching technique (Simpson and Oksanen, 2021) has been used to explore whether the modern (most recent) pollen samples from each site represent novel ecosystems. At the 10% cut-off level for defining close analogues, Alport Moor has modern analogue equivalents at ∼122 BC and ∼724 BC (Fig. 3f). Coldharbour Moor only has one close analogue, which is the pollen sample directly below the most recent sample, indicating that since the 17^th^ to 19^th^ century the record has represented a novel ecosystem. Bar Brook has a matching modern analogue at 18 cm depth (∼AD 1877), which mirrors the lower fire activity in the modern sample. Cranberry Bed, Emlin Dike and Withens Moor have no close modern analogues implying that the most recent samples may represent novel ecosystems.

## Discussion

### 1) Do fire and vegetation responses vary under different management practices?

Through assessing patterns of regeneration following fire, we identify that trends differ according to management and environmental setting and some sites may be more vulnerable to future fire activity, such as those already dominated by grasses with lower vegetation diversity (e.g. Withens Moor). Similar relationships have been identified in modern experiments where burn severity and pre-fire species composition were key factors determining resilience to fire (Davies et al., 2023; Baker et al., 2025). The CCA ordination plots (Fig. 5) reflect the relationships between pollen assemblage change and environmental variables at the six sites. Some relationships differ between management types with dissimilar patterns between vegetation, fire, pollen (Shannon) diversity and SCPs at different sites. In the CCA plot for Bar Brook, charcoal influx is associated with heathland pollen types (*Calluna vulgaris* and *Ericaceae*) and ribwort plantain (*Plantago lanceolata*), an important wider landscape grazing indicator. At Withens Moor, charcoal is most strongly associated with agricultural pollen indicators. At some sites, higher fire activity is associated with heathland (*Ericaceae*) and grasses while lower fire activity is associated with trees and *Sphagnum* moss. This relationship is consistent in different management areas with trees declining during periods of increased fire, and fires leading to promotion of heathland and grasses. Some participants in this study shared observations that prescribed burning has mixed ecological effects with individuals recognising its use as a tool for controlling vegetation and encouraging biodiversity in specific contexts, although others mentioned that burning can dry peatlands, increase fire-prone vegetation, and reduce *Sphagnum* cover. Participants also recognised that grazing intensity alters fire risk and vegetation composition. Past research has demonstrated that heavy grazing can reduce vegetation biomass, potentially reducing immediate fire risk (Smit and Coetsee, 2019), but such intense grazing can degrade ecosystems over time, and make recovery from fire slower (Davies, 2016; Davies et al. 2023).

**Figure 5.**
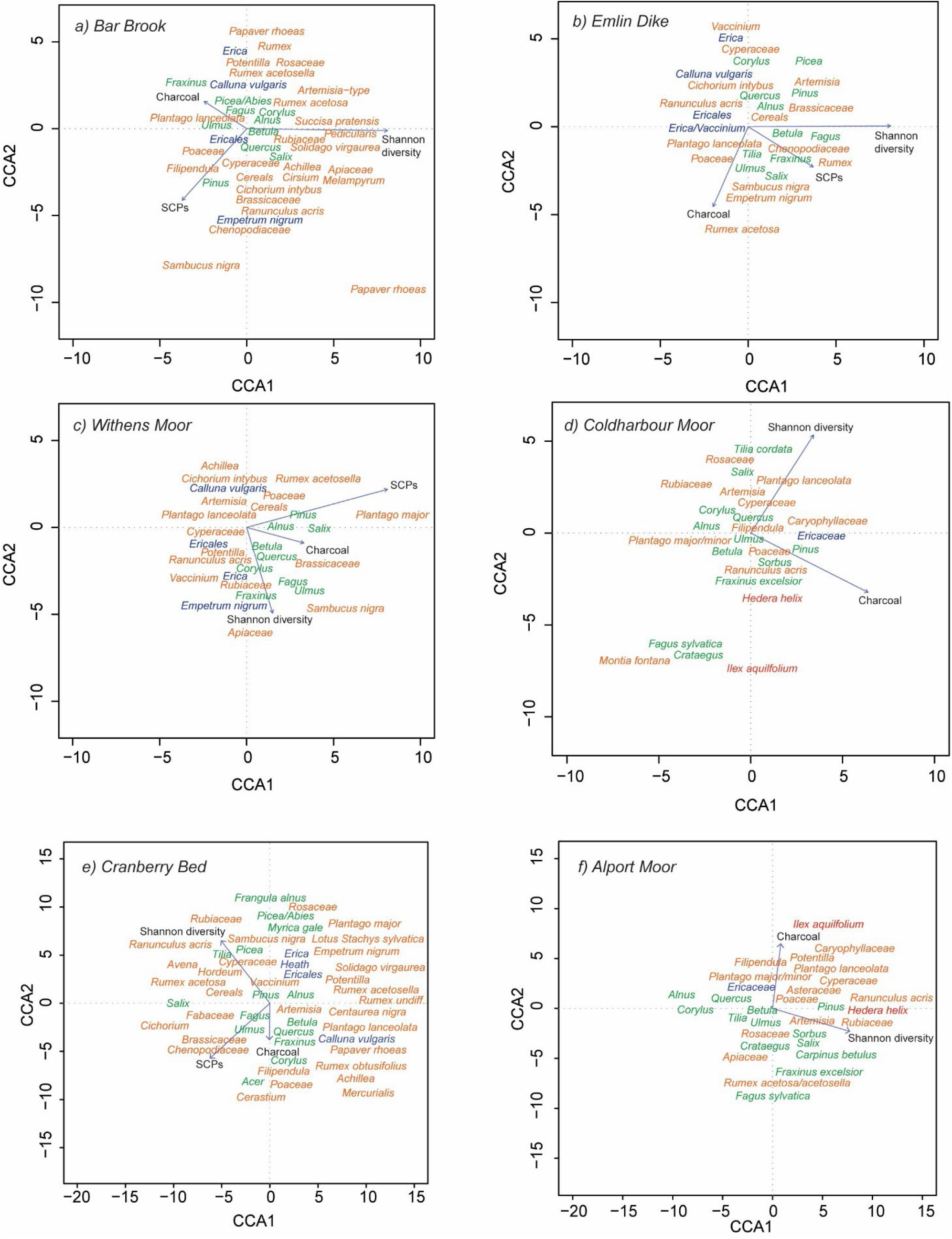
Pollen sample CCA (Canonical Correspondence Analysis) ordination plots: a) Bar Brook, b) Emlin Dike, c) Withen’s Moor, d) Coldharbour Moor, e) Cranberry Bed and f) Alport Moor. Green: trees, blue: heathland, and orange: herbs.

Stakeholder interviews revealed that pollution following fire events and fire impacts on biodiversity and environmental quality are major concerns. Patterns in the PCA ordination plots (Fig. 6) indicate that fire intensity in more recent times is not associated with the same species combinations as similar fire activity levels during earlier periods, implying that the relationship between fire and vegetation has shifted. The participatory research captured how different management practices reflect diverse social and cultural values, which encompass legacies of historical land-use. Different management approaches influence not only fire activity and vegetation responses, but also how communities interact with landscapes, shaping ecological outcomes. Conservation-focused areas often emphasise rewetting, biodiversity recovery, and limiting fire use. Whereas grouse moor management tends to use regular burning to maintain heather and reduce fuel, potentially lowering wildfire risk in the short term but with long-term ecological trade-offs. Participants indicated that integrated and adaptive management is most effective where knowledge-driven strategies using monitoring and local expertise are prioritised. This expertise needs to be integrated with scientific knowledge on the complex and dynamic relationships between fire, landscapes, climate and land use. Davies et al. (2016) emphasise that current debates often oversimplify burning despite its nuanced ecological role.

**Figure 6.**
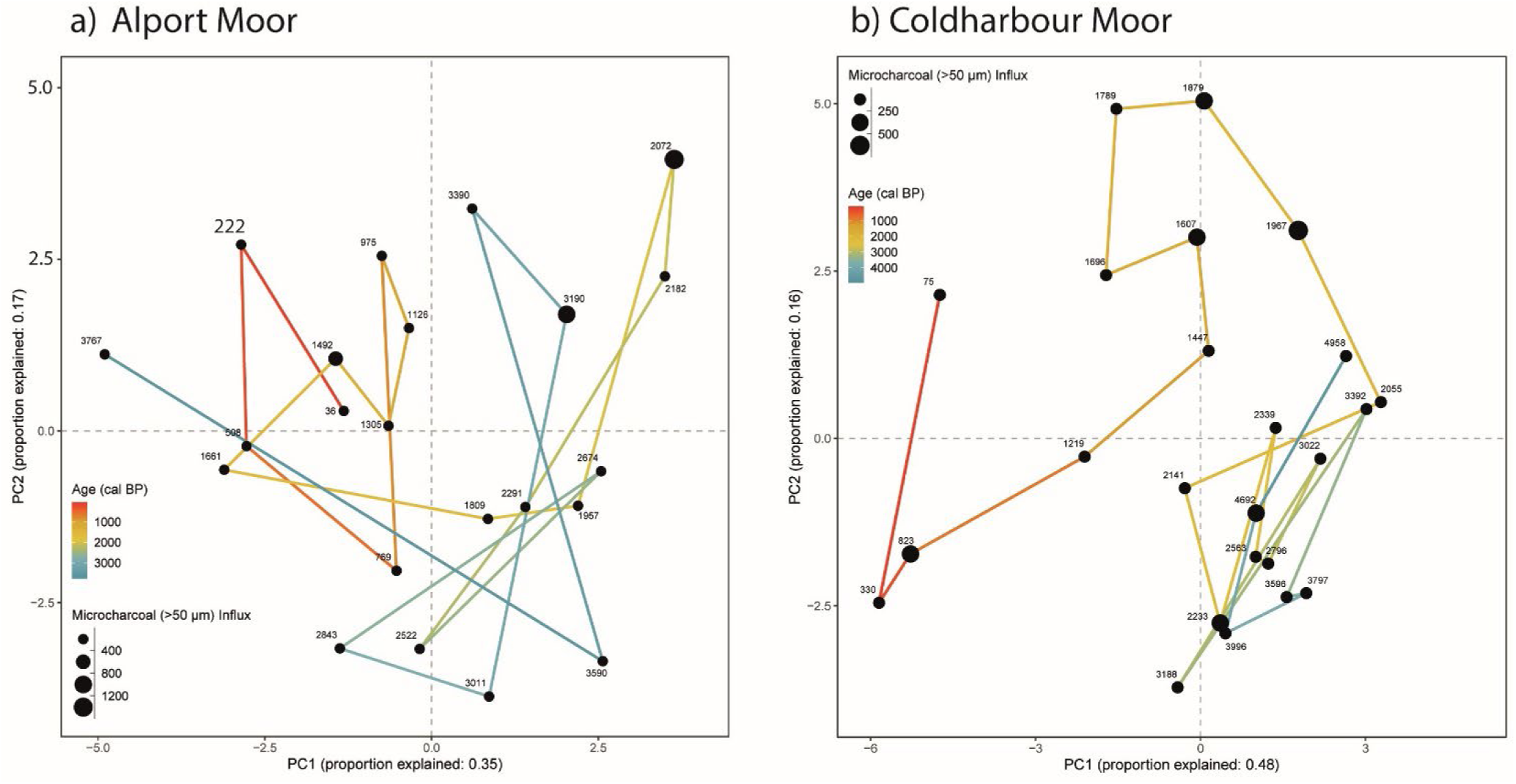
a) Alport Moor and b) Coldharbour Moor Principal Components Analysis (PCA) showing pollen sample similarity through time (blue: oldest to red: youngest). Size of sample points represents fire activity (charcoal concentration).

Restoration and management actions often include rewetting along with approaches to reduce wildfire risks and the frequency of burning. The participants in this research observe that rewetting encourages the return of *Sphagnum* moss. Degraded peatlands have been demonstrated to be less resilient to climate change (Loisel and Gallego-Sala, 2022) while systems that retain their hydrological conditions (waterlogged soils), peat-building processes, vegetation, biodiversity, and carbon-storage function are often more resilient. Earlier periods of the historic ecological records, when peatlands were less degraded, appear associated with more resilient ecosystems. Policy discussions have focussed on the regulation of burning practices, such as stricter constraints on controlled burning (UK Government, 2021), however, a sustainable approach to fire and land management needs to consider ecological health, wildfire reduction and the priorities of stakeholders (Colombaroli et al., 2019) and integrate public values into social-ecological system governance (Williams et al., 2020). Historic ecological data reveal that current management pressures are pushing some moorlands beyond their mid-to-late Holocene range of variability. ‘Shifting baseline syndrome’ (Soga and Gaston, 2018) may lead to assumptions that the modern conditions of the four records (Emlin Dike, Cranberry Bed, Withens Moor and Bar Brook) covering recent centuries are typical of these ecosystems. However, the earlier records from Coldharbour Moor and Alport Moor reveal that fire activity was lower, tree cover and heathland were more abundant, and grass dominance was lower during prior centuries and millennia, which concurs with other Holocene scale palaeoecological research in the PDNP (Tallis, 1991; Taylor et al., 1994; Long et al., 1998; Simmons et al., 2022).

### 2) How have fire and burning affected landscapes and ecosystems historically in the PDNP?

The historical ecological evidence, showing land-use change over time (e.g. grazing, burning, and heather management), aligns with points made by interviewees in this study focused on the unique character of the PDNP. Historic ecological data reveal how coexisting and layered land management legacies over recent centuries shape the current landscape mosaic. This uniqueness is captured in the site-specific records, which allow exploration of when and how ecological complexity emerged. The records reveal that fire has continuously shaped the PDNP landscape with varying impacts on vegetation composition. Some sites show a shift from heathland to grass dominance after persistent and increased fire activity (e.g. Withens Moor and Cranberry Bed), while others show recovery of heathland species as fire activity decreases (e.g. Emlin Dike, Coldharbour Moor, Alport Moor and Bar Brook). Magnan et al. (2012) found that fire temporarily altered the composition of *Sphagnum* mosses and other peatland vegetation, but these changes were generally limited to a few decades, after which vegetation revered back to its pre-fire state. They found that ombrotrophic peatlands show high resilience to surface fires, with only transient vegetation shifts and minimal long-term impact on carbon accumulation. This implies that some peatland types are more resilient than others, as identified by Mauquoy and Yeloff (2007), who demonstrated that some bogs showed long-term stability despite environmental shifts, while others were more vulnerable to change.

Modern ecosystems in some areas differ from historic vegetation assemblages and may require different types of restoration or management. Sites like Emlin Dike, where rotational burning is used for grouse moor management, show different ecological patterns and responses to fire compared to more degraded areas like Withens Moor. Shifting land management practices, including peatland drainage and prescribed burning of moorlands (Ramchunder et al., 2009; Davies, 2010; Fyfe et al., 2012), coincide with widespread peatland drying (Swindles et al., 2019). Other research concludes that human use of fire may have now exceeded the ability of peatlands to resist burning (Davies et al., 2016; Sim et al., 2023). The interview participants have also observed that landscape drying has made some deep peat areas especially flammable. Open moorland has dominated the Peak District for over 2000 years while fragmented tree and shrub cover persisted until the 17th century. Grazing led to heather moorland replacing tree cover, and fire, when used in management, contributed to the contraction of trees and shrubs. In recent centuries, pollution (evidenced by SCP data), burning, and grazing have reshaped vegetation communities (Davies, 2016).

The records from Coldharbour and Alport Moor illustrate that lower intensity fire was a persistent landscape feature in the millennia prior to the industrial period. These records show different relationships between vegetation and fire over longer timescales (i.e. the last 3000 years) compared to the more recent records over the last few hundred years, as trees and heath were abundant in the wider landscape despite regular fire occurrence. The historic ecological data confirm that long-term fire activity has cumulative consequences while modern studies show that higher frequency blanket peat burning over longer periods significantly reduces carbon sequestration (Garnett et al., 2000). Kuhry (1994) found that fires have limited short-term ecological effects but can drastically slow long-term carbon storage in boreal peatlands if they occur more often. Climate also plays a significant role in peatland resilience to fire (Loisel and Gallego-Sala, 2022). We infer that the historic ecological records appear to show decreased fire activity during the Little Ice Age climatic episode (14^th^ - 19^th^ centuries AD). The Medieval Warm Period (MWP) (Medieval Climatic Anomaly) (9^th^ - 13^th^ centuries AD) involved warmer conditions in northwest Europe, allowed more land to be used for cultivation, and saw increased burning in peatlands (Olsson et al., 2009; Costello et al., 2021). The MWP can provide a “natural experiment” to explore the consequences of warmer climatic conditions for an extended period alongside the increasing use of burning as a land management tool (Marlon et al., 2013; Turner et al., 2014; Costello et al., 2021). The PDNP landscape appears to have remained vegetationally diverse throughout the MWP despite the persistent occurrence of fire, therefore the level of fire activity and land management during this time appears sustainable in terms of maintaining ecosystem resilience. Although regionally variable, MWP temperatures are estimated to have been similar to the period AD 1960-1990 (Goosse et al., 2006), however, current and future climate trends exceed this threshold with the last decade the warmest in at least the past 1000 years alongside increasing drought events (IPCC, 2022), which could be associated with more peatland fires in the future.

The participatory research in this study highlights the legacy impacts of historic wildfire and burning with topics, such as ecological impacts, land-use change, biodiversity change, soil erosion, flooding, and displacement of wildlife, highlighted by stakeholders. Periods of low fire frequency have reduced investment in preparedness, leading to fuel accumulation and larger, more destructive fires later. Interview participants emphasised a need to better understand long-term ecological responses, including how different management regimes influence post-fire regeneration and ecosystem recovery. Some participants appreciated the value in understanding historical ecosystem responses to fire, noting that examining what vegetation existed before and how fire interacted with vegetation helps identify thresholds and tipping points. Other interviewees described how 19th- and 20th-century practices like draining peatlands, intensive grazing, and frequent burning may have made modern landscapes more fire-prone.

### 3) How can long-term historic ecological records inform wildfire management?

Prior vegetation condition, site hydrology, and previous land use are factors that play significant roles in a peatland’s susceptibility to and recovery from fire (Davies et al., 2023). Moss (*Sphagnum*) and sedge (*Carex*) were more abundant in the past as evidenced by the longer palaeo records from Alport and Coldharbour Moor, therefore earlier periods in these records represent conditions that should need to be promoted in restoration efforts (Noble et al., 2018). Points in the past with higher landscape heterogeneity and favourable ecological conditions can be used as baseline targets for restoration (Guilfoyle et al., 2024), but trends and relationships can be highly variable between sites (Chambers et al., 2013). Intensive land use, such as over-grazing or regular controlled burning for agriculture or grouse management, can reduce *Sphagnum* moss cover, which is essential for peat formation, and promote heath dominance, therefore these management strategies need careful consideration. Coldharbour Moor is an example of a site where burning has promoted heath cover, which has exceeded the historic baseline in the most recent part of the record, whereas Bar Brook is the most ecologically favourable site with abundant *Sphagnum* moss suggesting that conservation has been successful here. However, the palaeoecological datasets do not differentiate between different *Sphagnum* species, therefore further research into the specific species and whether these are peat forming types is needed, for example, using plant macrofossil data to capture local vegetation composition (McCarroll et al., 2016). Restoring waterlogged conditions, encouraging *Sphagnum* regeneration, and actions that encourage vegetation diversity, will help support healthier ecosystems that are more resilient to fire. The record from Emlin Dike represents a resilient ecosystem, while Withens Moor characterises a site that has lost resilience. Cranberry Bed and Bar Brook are beginning to show a similar pattern to Emlin Dike with an increase in heathland and trees following declining fire and pollution (SCPs), but the sites require regular monitoring to ensure that this pattern is maintained. Alport Moor has been managed by rewetting, gully blocking and revegetating with cotton grass (E*riophorum*) and heather (*Ericaceae*). The historic ecological record shows that grass dominance was not typical here and native woodland and *Sphagnum* moss were previously abundant in the area. The site may return to previous conditions with suitable management, and decreased burning may support the reestablishment of *Sphagnum* moss, which is also supported by prior research (Magnan et al., 2012; Milner et al., 2021). Issues surrounding fuel load accumulation increasing risk for wildfire were frequently mentioned by participants in this research. Cutting and removal of vegetation to manage biomass and fuel build up could be a viable option instead of burning, however, participants highlighted that cutting is not suitable everywhere and its effectiveness varies with terrain. Further, managed grazing can also help to manage fuel load, as demonstrated by Smit and Coetsee (2019).

While the term "palaeoecology" was rarely used explicitly, interviewees in this study consistently valued long-term perspectives and fire history. There is clear appreciation for using the past to inform future wildfire resilience and land restoration strategies, such as through defining restoration goals based on past conditions and identifying the drivers of environmental change. Sites with known historical plant communities, such as wetter or more biodiverse assemblages, can provide models for restoration goals that reduce fire susceptibility. Interviewees implied that long-term records are most useful when combined with traditional knowledge, land user experience, and ecological monitoring. Wildfire risk reduction requires vegetation management, and the promotion of fire breaks, community awareness, appropriate regulations, land-use planning, and funding and resources for risk management. Wildfire risk needs to be included in climate resilience plans alongside multiple organisations working collaboratively to share best practices and resources (Otero et al., 2018), which is already in place with initiatives such as the Moors for the Future Partnership and the PDNP Fire Operations Group. The results of this study support ideas around careful consideration of controlled burning on peatlands and highlight that diverse vegetation types maintained by wetter soils are more resilient to fire and can be used to support natural regeneration. The palaeo records reveal that woodlands were more extensive and abundant in the past PDNP landscape, which illustrates the potential of restored native woodlands to create natural firebreaks. In support of this, the IPCC (2022) highlight that restoring less flammable native oak (*Quercus*) forest communities would reduce fire risk and erosion compared to recent forest plantations involving flammable trees (e.g. pine).

The changing relationships between vegetation, land use and fire over long time periods illustrate that a return to “natural” conditions may not necessarily be realistic, so cannot be the key priority for restoration projects in diverse multi-use fire-adapted landscapes. Addressing environmental challenges will not involve simply reducing burning, rewetting and revegetating with typical moorland species, as the vegetation-fire relationships that have developed over multiple millennia are complex. Individual sites will require careful and tailored management that learns from historic legacies and recognises fire as an integral component of the historic and current landscape. Various policies have been developed and actions outlined in recent years that aim to protect peatlands and support sustainable fire management (Glaves, 2020). Historic ecological data could be useful in assessing the suitability of policies and informing their implementation.

## Conclusion

The historical ecological records from the PDNP offer insights into the relationships between fire, vegetation, climate, and land management over recent centuries to millennia. Sites like Emlin Dike demonstrate that ecosystems can recover and maintain resilience when fire activity changes, while sites such as Withens Moor, which has become dominated by purple moor grass and has lost resilience, highlight the challenges of restoring landscapes that have reached ecological thresholds. This research highlights the importance of understanding socio-ecological systems and adopting integrated wildfire management strategies that account for ecological, social, and cultural aspects. Participants provided insights into the complexities of managing diverse peatlands in relation to changing fire dynamics. Persistent fire activity and intensive land use has altered the relationship between vegetation and fire, indicating a significant role for future effective management. Lower intensity fire activity in the earlier periods of the palaeoecological records, alongside more intact peatland function, supported greater landscape heterogeneity and resilience. Restoring peatlands through rewetting, promoting *Sphagnum* moss growth, and limiting controlled burning, are key measures to enhance resilience to future fire. Re-establishing native woodland and improving hydrological conditions are also positive actions for reducing wildfire risk, alongside continued monitoring or the consequences of fire and conservation or restoration actions.

## Supporting information

Supplementary Information: Woodbridge et al.

## Author contributions

Jessie Woodbridge, Gina Kallis, Laura Scoble, Francis Rowney, Claire Kelly and Althea Davies were central to the conception and design, acquisition of data, analysis and interpretation of the data. JW drafted the manuscript and GK, LS, FR, CK and AD revised the content critically contributing intellectually. LS carried out laboratory analyses and FR carried out some of the data analyses. AD contributed pollen data for analysis and commented on the manuscript providing critical feedback. All authors contributed critically to the drafts and gave final approval for publication.

## Acknowledgments

This research has been carried out as part of the “PeakfireRes” project funded by the Royal Geographical Society and Association for Environmental Archaeology. Radiocarbon-dating was also supported by funding from the NERC National Environmental Isotope Facility. We are grateful to many individuals who generously contributed their time to share their expertise and discuss their experiences. We also thank anonymous reviewers who provided useful comments and made valuable suggestions to improve this publication.

## Data availability

The data will be made available in Neotoma upon publication. Neotoma Database: https://www.neotomadb.org/. R scripts will be made available by the authors upon request.

## Funding details

This research was funded by the Royal Geographical Society, Association for Environmental Archaeology and NERC National Environmental Isotope Facility.

## Disclosure statement

The authors report there are no competing interests to declare.

## Biographical note

Jessie Woodbridge is a Lecturer in Ecosystem Resilience who researches past ecological change and human-environment interactions, and links this to modern environmental challenges. Gina Kallis is a multi-disciplinary social science Research Fellow and Social Research Consultant. Laura Scoble is a researcher in physical geography, specifically focusing on palaeoecology, environmental change and applied chemistry. Claire Kelly is an environmental social scientist. Her work focuses on understanding and evaluating the relationships between society, the environment and the economy using interdisciplinary co-design and participatory research methods. Francis Rowney’s background is in palaeoecological research, and he is now a practitioner in restoration and conservation. Althea Davies is a historical palaeoecologist whose work examines long-term environmental change, landscape history, and the interactions between people and past environments.

## Notes

### Competing Interest Statement

The authors have declared no competing interest.

