## Supplementary Information: Woodbridge et al. for "Historical ecology and stakeholder perspectives can inform peatland fire management"

**Supplementary Information: Woodbridge et al. Preprint**

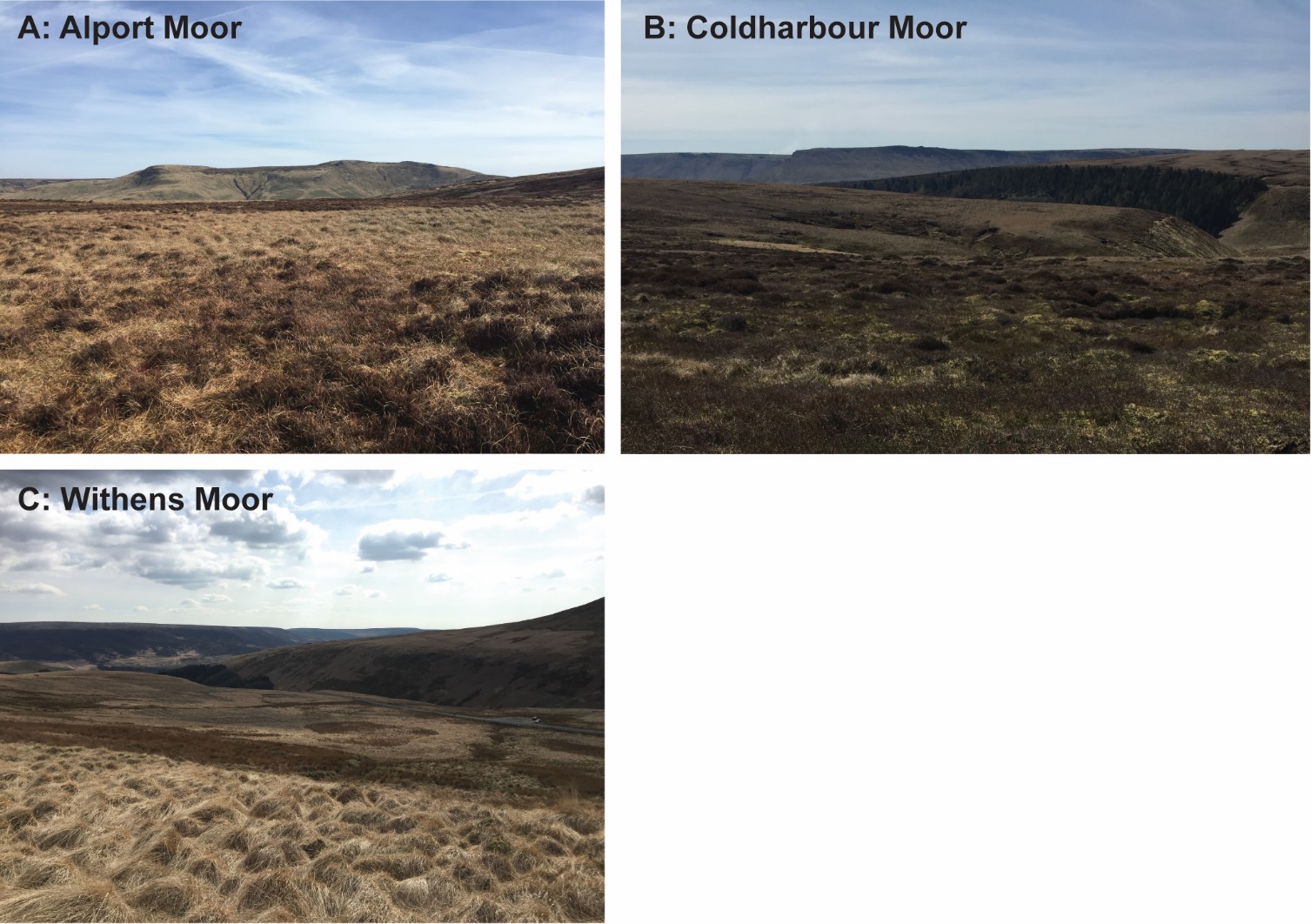

**Supplementary Information 1.** PDNP peat coring location photos - A: Alport Moor, B: Coldharbour Moor and C: Withens Moor.

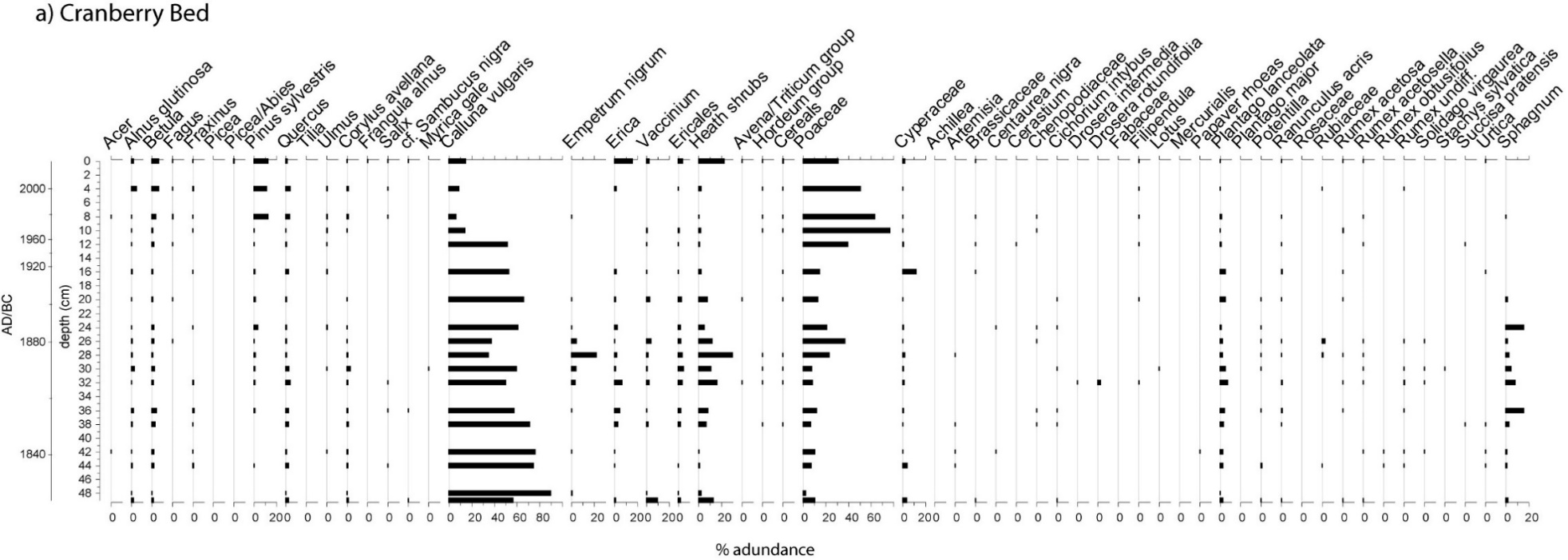
**Supplementary Information 2.** Pollen percentage diagrams for a) Cranberry Bed, b) Bar Brook, c) Emlin Dike, d) Withens Moor, e) Coldharbour Moor, and f) Alport Moor showing all land pollen taxa and *Sphagnum* moss.

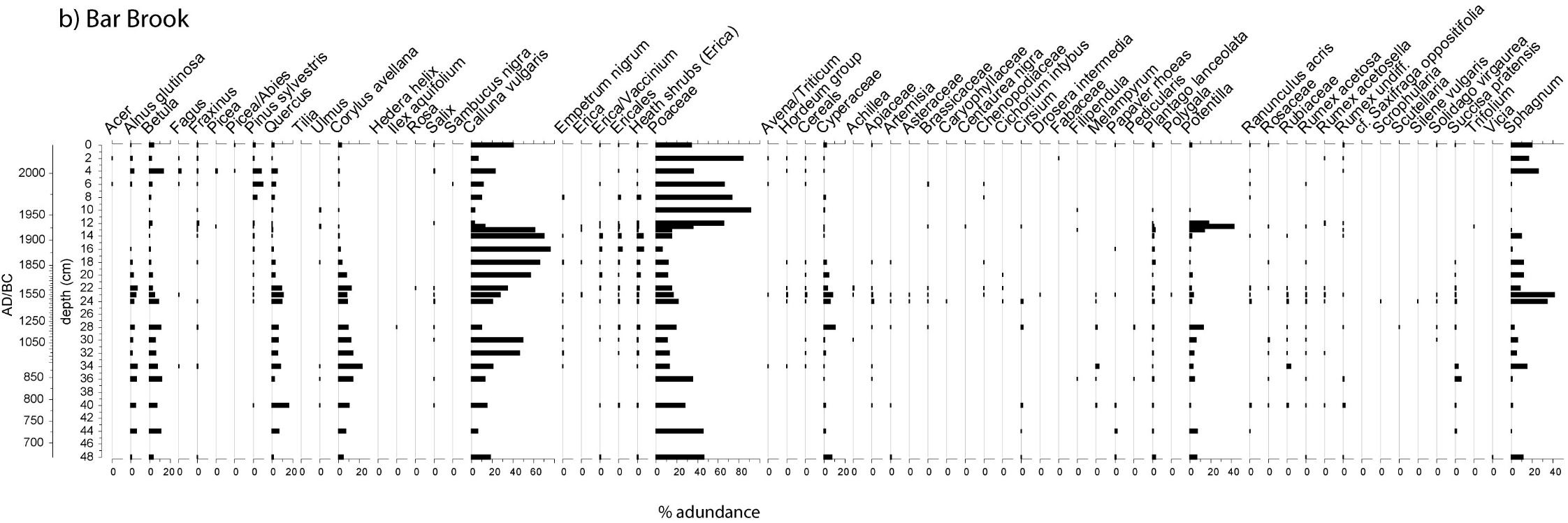

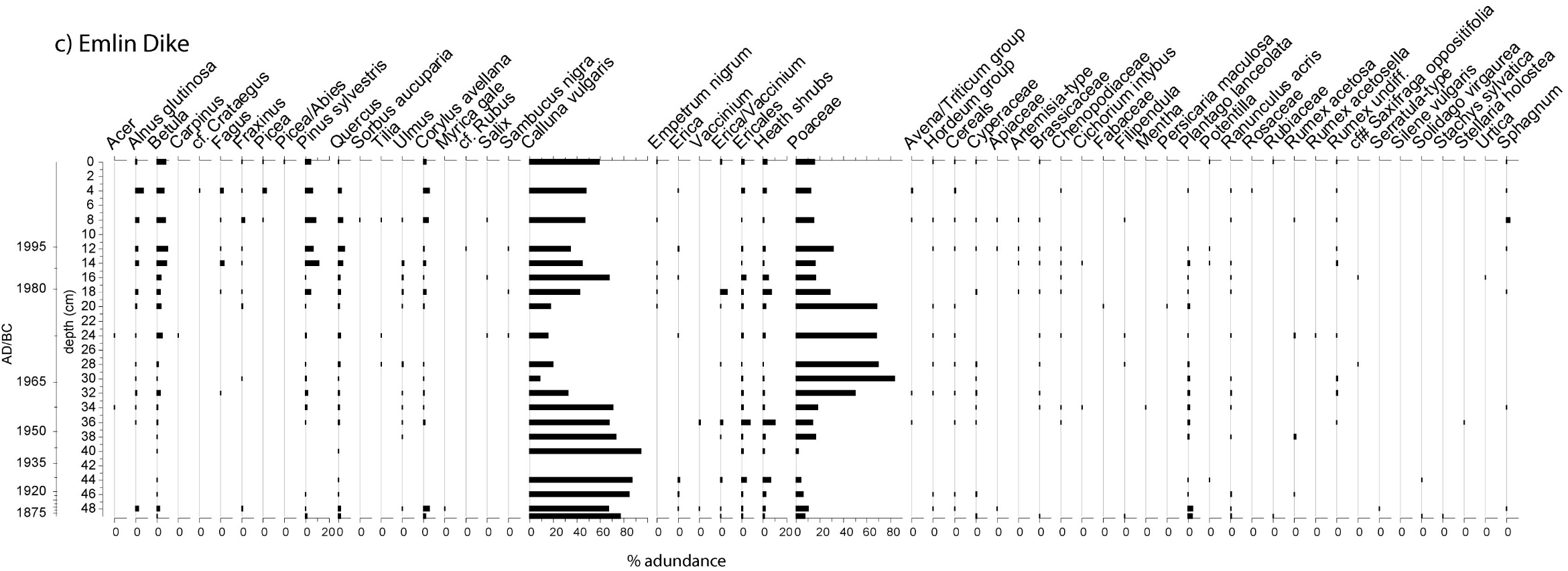

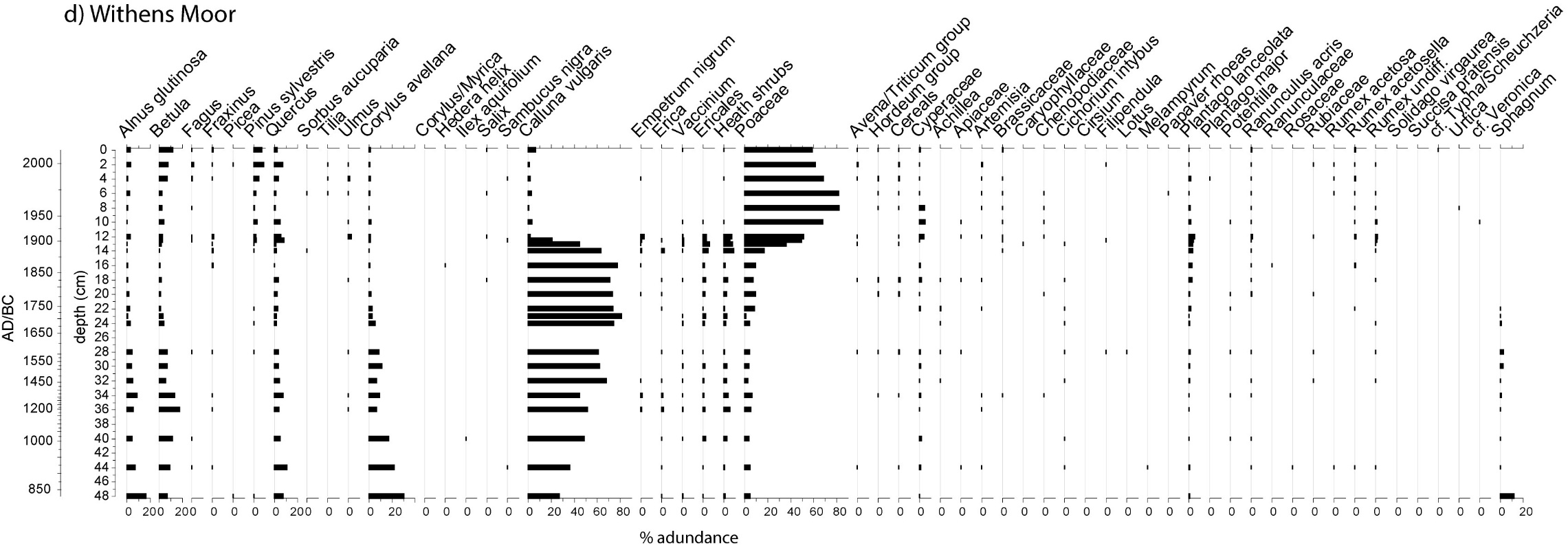

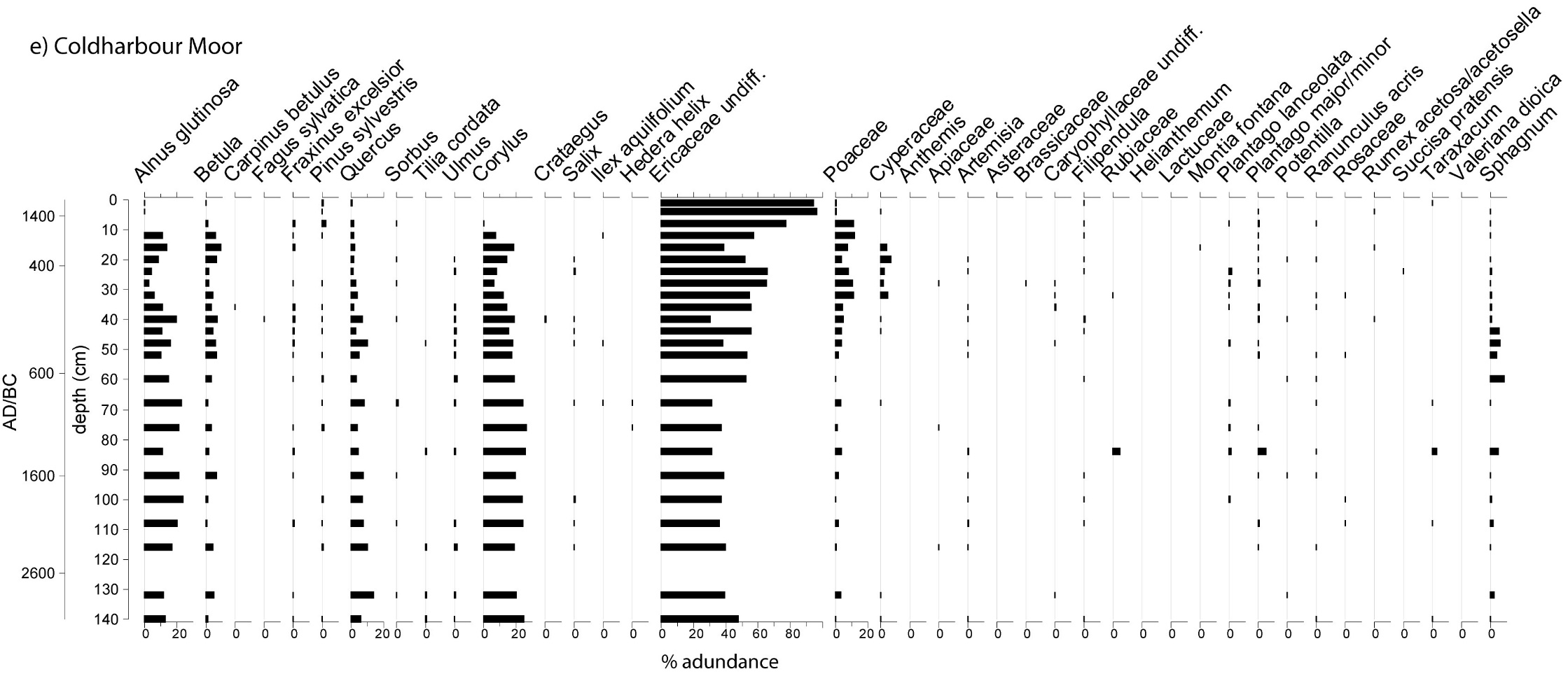

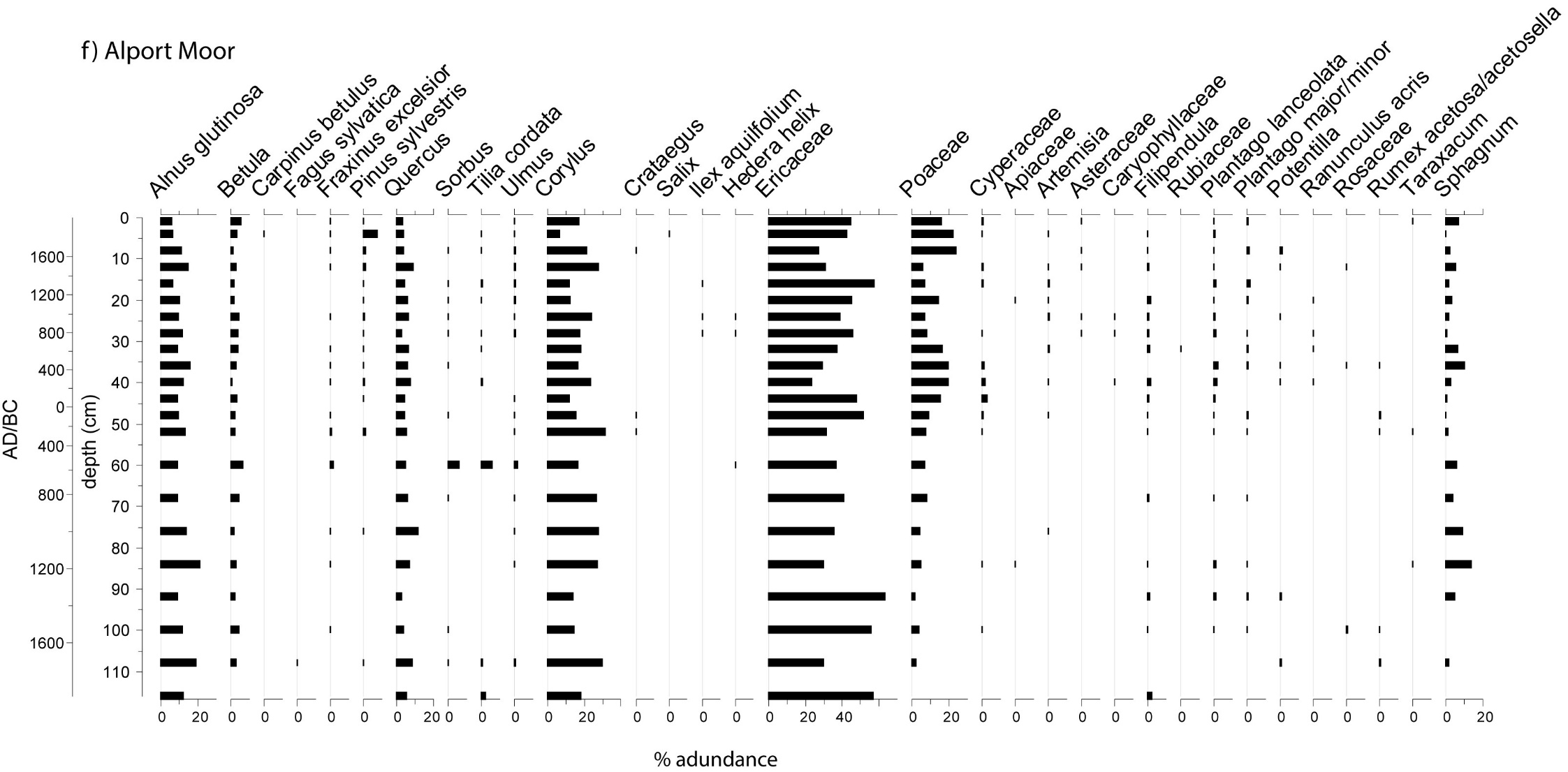

**Supplementary Information: Table 5** Spearman’s Rank correlations between the datasets from the six PDNP palaeoecological sites.

| **Spearman’s Rank** | **Alport Moor** | **Coldharbour Moor** | **Bar Brook** | **Cranberry Bed** | **Emlin Dike** | **Withens Moor** |
| --- | --- | --- | --- | --- | --- | --- |
| **Charcoal concentration/ influx** | Positive relationships with herbs/grazing indicators (e.g. plantain (*Plantago lanceolata*) (**r=0.476, p=0.025**).  Negative relationship with % trees (**r=-0.514, p=0.014**). | Negative relationships with hazel (*Corylus*) (**r=-0.473, p=0.02**). | Strong negative relationship with *Sphagnum* moss (**r=-0.664, p=0**), cereals (**r=-0.77, p=0**). Negative relationships with trees (alder: *Alnus*) (**r=-0.533, p=0.007**). | Negative relationships with vegetation evenness (**r=-0.505, p=0.033**). Negative relationship with alder (*Alnus*) (**r=-0.474, p=0.047**). | Negative relationship with several tree types: pine (*Pinus*) (**r=-0.519**), oak (*Quercus*) (**r=-0.457**) hazel (*Corylus*) (**r=-0.497**), heather (*Calluna vulgaris*) (**r=-0.495**, **p=0.027**). Positive relationship with grass (**r=0.556, p=0.011**) | Positive relationships hazel (*Corylus*) (**r=0.548, p=0.006**). |
| **Shannon diversity (pollen)** | Diversity positive correlated with herbs (**r=0.578, p=0.005**). Negative relationship with heath (*Ericaceae*) (**r=-0.695, p=0**)  No relationship between diversity and charcoal. | Negative relationships with trees: birch (*Betula*) (**r=-0.547, p=0.006**) and ash (*Salix*) (**r=-0.463, p=0.023**). | Positive relationships with herb and tree types (e.g. alder (*Alnus*) (**r=0.798, p=0**), oak (*Quercus*) (**r=0.831, p=0**), hazel (*Corylus*) (**r=0.777, p=0**). | Correlations with many taxa e.g. herbs (heather (*Calluna vulgaris*) (**r=-0.554**), heath (*Erica*) (**0.692**), cereals (**0.511**) and trees: alder (*Alnus*) (**r=0.595**), pine (*Pinus*) (**r=0.577**), oak (*Quercus*) (**r=0.552**). | Significant relationships with herbs (cereals: **r=0.541**) and trees: alder (*Alnus*) (**r=** **0.871**), oak (*Quercus*) (**r=0.785**), pine (*Pinus*) (**r=0.749**) hazel (*Corylus*) (**r=0.79**). | Positive relationship with tree types: alder (*Alnus*) (**r=0.437, p=0.033**), oak (*Quercus*) (**r=0.814, p=0**), hazel (*Corylus*) (**r=0.416, p=0.043**) and (cereals: **r=0.422, p=0.04**). |
| **SCPs (Spheroidal Carbonaceous Particles**  **Peat humification (climate proxy)** | No SCP data.  No significant relationships identified with humification data. | No SCP data.  Humification significantly correlated with number of pollen taxa (**r=-0.583, p=0.005**). | SCPs: Negative relationship with *Sphagnum* (**r=-0.415, p=0.043**) diversity (**r=-0.73, p=0**) and several pollen taxa. | SCPs: Negative relationship with *Sphagnum* (**r=-0.505, p=0.032**) and significant relationships with several pollen taxa. | SCPs: Negative relationship with heather (**r=-0.649, p=0.002**) and positive with grass (**r=0.588, p=0.006**) and other herbs. | SCPs: Negative correlation with *Sphagnum* (**r=-0.55, p=0.005**), heather (**r=-0.711, p=0)**, pine (**r=0.786, p=0)**, and grass (**0.831, p=0**). |

**Supplementary Information: Table 6** Quotes from individuals interviewed during participatory research, illustrating personal perspectives, experiences, and attitudes related to peatland management and restoration practices.

**Quotes from anonymous interview participants**

“*The wildlife, of course, because all these fires tend to be bang in the middle of bird nesting […] when you look at one we had this year, the wind was blowing that strong. I mean, we found mountain hares burnt and, you know, I mean, that's the fastest land mammal in the UK. And it can't be- if it couldn’t outrun a fire, then that must have been, yeah, it must have been shifting*.” (Interviewee)

"*Very, very dangerous, especially in the Peak District where the wind, the winds that sort of tend to fan a bad wildfire tend to come from the southeast or east. Between sort of Sheffield and Manchester, there's a huge block of maybe 12 miles of solid heather. That is going to go up and not only highly dangerous, and it releases all this carbon that we’ve been sequestering, it also releases a sort of pollution from the Industrial Revolution that we dropped on prevailing westerly winds from places like Manchester. So it’s a really big problem*.” (Interviewee)

*“There's a study on the Saddleworth Moor- the amount of metals that it released and pollutants that it released in watercourses […] It obviously releases carbon, it releases sediment into water courses. So there's a whole range of scientific- environmental impacts […] And obviously, that bleeds into socio-economic ones then because of costs of cleaning water, flooding that occurs from denuded moorlands, some point down the line.”* (Interviewee)

*“I think to have that security, if you like, that if we do have one [wildfire], it’s one phone-call into the FOG partnership and I know that we could get access to specialist support from any one of those partners, it’s a massive benefit and vice versa*”. (Interviewee)

*“In terms of economic loss, an individual was paying 25,000 pounds a day for the helicopter, come in, pick up water from the reservoir and dump it on hotspots*.” (Interviewee)

“*The immediate economic return from the land in terms of sheep grazing or grouse shooting, that also goes down. The need for restoration increases so the cost input goes up. So those are immediate very local impacts*…” (Interviewee)

“*We have a unique habitat and unique land use, multiple land use, you know, we've got grazers, landowners, sporting rights, mineral rights, they’re all layered on top of each other all on one patch of land. So it's highly complex, and unique*.” (Interviewee)

“*Quite often we'll have a policy that comes down the line from government and unless it's met with people's experience from the ground up, quite often misses the mark, and you get unintended consequences.”* (Interviewee)

*“Trying to put together the practitioner with the scientist is really important, and the scientists must have, you know, due regard, respect for the view of the practitioner because otherwise, you know, we're in for- you only need to look at things like the Staybridge fire to realise something is amiss.”* (Interviewee)
